# An open, fully-processed data resource for studying mood and sleep variability in the developing brain

**DOI:** 10.1101/2025.05.01.651544

**Authors:** Juliette B. H. Brook, Taylor Salo, Audrey C. Luo, Joëlle Bagautdinova, Sage Rush, Aaron F. Alexander-Bloch, Erica B. Baller, Monica E. Calkins, Matt Cieslak, Elena C. Cooper, John A. Detre, Mark A. Elliot, Damien A. Fair, Phoebe Freedman, Philip R. Gehrman, Ruben C. Gur, Raquel E. Gur, Arno Klein, Nina Laney, Timothy O. Laumann, Kahini Mehta, Kathleen Merikangas, Michael P. Milham, Jonathan A. Mitchell, Tyler M. Moore, Steven M. Nelson, Kosha Ruparel, Brooke L. Sevchik, Sheila Shanmugan, Haochang Shou, Manuel Taso, Lauren K. White, Daniel H. Wolf, M. Dylan Tisdall, David R. Roalf, Theodore D. Satterthwaite

## Abstract

Brain development during adolescence and early adulthood coincides with shifts in emotion regulation and sleep. Despite this, few existing datasets simultaneously characterize affective dynamics, sleep variation, and multimodal measures of brain development. Here, we describe the study protocol and initial release (n = 10) of an open data resource of neuroimaging paired with densely sampled behavioral measures in adolescents and young adults. All participants complete multi-echo functional MRI, compressed-sensing diffusion MRI, and advanced arterial spin-labeled MRI. Behavioral measures include ecological momentary assessment, actigraphy, extensive cognitive assessments, and detailed clinical phenotyping focused on emotion regulation. Raw and processed data are openly available without a data use agreement and will be regularly updated as accrual continues. Together, this resource will accelerate research on the links between mood, sleep, and brain development.

## Introduction

Affective instability is characterized by frequent mood shifts and disturbances in emotional intensity;^1^ affective instability is related to similar constructs such as mood instability and emotional dysregulation.^2^ Affective instability is present in multiple psychiatric disorders, including borderline personality disorder (BPD), attention-deficit hyperactivity disorder (ADHD), post-traumatic stress disorder (PTSD), and major depressive disorder (MDD).^3^ Typically emerging when youth learn to self-regulate changes in their affective states or mood,^1^ affective instability has been associated with suicidality and suicide attempts.^4^ Despite increasing rates of mental disorders and suicide in youth,^5^ the neurodevelopmental underpinnings of affective instability remain sparsely studied and poorly understood.^6^ Here, we describe the study protocol and first data release of a new open data resource focused on how variation in brain development may relate to affective instability in youth.

Affective instability is challenging to measure. While there are numerous self-report scales (e.g., Affective Control Scale,^7^ Difficulties in Emotion Regulation Scale,^8^ etc.) and semi-structured interviews (e.g., Emotion Regulation Interview,^9^ Structured Clinical Interview for DSM-V Axis II Personality Disorders,^10^ etc.) that assess aspects of affective instability, their retrospective nature limits accuracy and can be susceptible to memory distortion.^11^ Ecological momentary assessment (EMA) offers a promising alternative by measuring participants’ self-reported symptoms and activities as they occur in daily life, providing contextual richness and naturalistic data.^12^ These strengths have led to increasing use of EMA in studies of BPD,^13,14^ MDD,^15^ and other related disorders in childhood.^16^

Sleep quality is closely tied to affective dynamics and emotion regulation in youth. Substantial evidence indicates that sleep disruption (e.g., sleep deprivation, insomnia) drives emotion dysregulation and predicts the onset of mood disorders in adolescence.^17–22^ Additionally, sleep is thought to be linked to critical plasticity mechanisms of healthy brain development,^23–26^ with impaired sleep contributing to variation in brain structure and function.^27–33^ Daytime physical activity has also been associated with affective dynamics, where lower levels of physical activity tend to worsen emotion regulation and mood.^34,35^ While sleep and physical activity are often assessed with parent or self-report questionnaires, research-grade actigraphy recordings provide sensor-based data that overcome questionnaire-related limitations of recall bias. Actigraphy tracks sleep and physical activity for up to several weeks in real-world settings, providing a unique opportunity to collect more accurate, dense, and ecologically valid measures of a person’s lifestyle.^36–38^

Neuroimaging data can be integrated with the high-quality behavioral measures described above to further characterize the neurodevelopmental underpinnings of affective dynamics. However, recent evidence suggests that due to small effect sizes, large samples are necessary to uncover generalizable brain-behavior relationships, posing a significant methodological challenge to understanding how variation in brain development connects to affective instability and mood dysregulation.^39,40^ Small effect sizes observed may be due, in part, to poor reliability and inaccurate measurements of key behavioral phenotypes,^39,41,42^ un-modeled individual variation in neuroimaging measures,^43,44^ and biological heterogeneity.^45–47^ However, studies can improve signal and minimize noise by collecting precise individualized measures of behavioral and brain data.^44^ The recent advent of multi-echo functional MRI (ME-fMRI)^48,49^ and other enhanced imaging sequences (like compressed sensing diffusion spectrum imaging; CS-DSI)^50^ may allow for identification of person-specific variation with less data.^48^ In particular, multi-echo fMRI facilitates accurate characterization of personalized functional networks (PFNs). PFNs delineated using ME-fMRI hold significant promise in improving the effect size and generalizability of translational studies.^48^ Thus, pairing contemporary EMA and actigraphy procedures that densely sample mood and sleep with more precise imaging methods may hold promise for linking variation in affective dynamics to brain development in youth.

In this paper, we describe the protocol and initial data release from a new study that leverages recent advances in characterizing complex behavior and non-invasive brain imaging to investigate affective instability in youth. Specifically, we employ a combination of EMA techniques and specialized self-report scales to assess affective instability and emotion regulation. Furthermore, the study measures sleep via wearable actigraphy devices in addition to EMA and other self-report measures. Perhaps most importantly, the study includes advanced imaging methods – such as ME-fMRI, CS-DSI, and advanced arterial spin labeled (ASL) perfusion MRI – to characterize structural and functional brain development. Notably, both raw and processed data are fully de-identified and openly shared without a data use agreement. We anticipate that this data resource will accelerate translational research on affective instability in youth.

## Methods

### Participants

This study aims to recruit 100 individuals between the ages of 13 and 23. Participants must be proficient in English, able to understand the study procedures, and agree to participate by giving written informed consent or assent. Exclusion criteria include any significant medical or neurological illness that may impact brain function or a history of pervasive developmental disorder, psychosis, bipolar disorder, or clinically significant current substance misuse. Furthermore, participants who present with acute intoxication with alcohol or other substances or lack a mobile device with capabilities to complete the study procedures are excluded. Additional exclusion criteria include pregnancy, implanted ferrous metal, claustrophobia, and other contraindications to MRI. Initial recruitment was focused on young adults as study procedures were being finalized; the initial data release includes 10 participants (mean age = 21.5, SD = 1.33; M:F 4:6). Recruitment is ongoing to reach the full sample size of 100 participants.

Participants are primarily recruited through medical record review at the Children’s Hospital of Pennsylvania (CHOP) and Penn Medicine. In addition, individuals who had previously participated in research within the Department of Psychiatry may be recontacted to participate in the current study. Other recruitment strategies include physician referrals within the Penn Medicine and CHOP systems, announcements on the lab website, and advertisements around Philadelphia. All study procedures are approved by the Institutional Review Boards of both the University of Pennsylvania and CHOP; all participants provide written informed consent. For participants under the age of 18, a parent or legal guardian provides informed consent and the minor provides informed assent.

### Behavioral and Clinical Assessments

This study includes an array of assessments. These include self-report measures, clinical assessments, a cognitive battery, mobile phenotyping with ecological momentary assessments (EMA), actigraphy via a wearable wristwatch-like device, and MRI. **Figure 1** presents the study workflow of assessments. First, the study team identifies and recruits participants through a phone call screener. Second, participants complete the in-person imaging visit at the Hospital of the University of Pennsylvania. Participants also begin the EMA procedures and actigraphy procedures following the in-person imaging visit. Participants are provided the actigraphy device fully charged as well as a prepaid shipping envelope to return the device after three weeks. Finally, participants complete a remote clinical assessment session in the days following the imaging visit.

**Figure 1.**
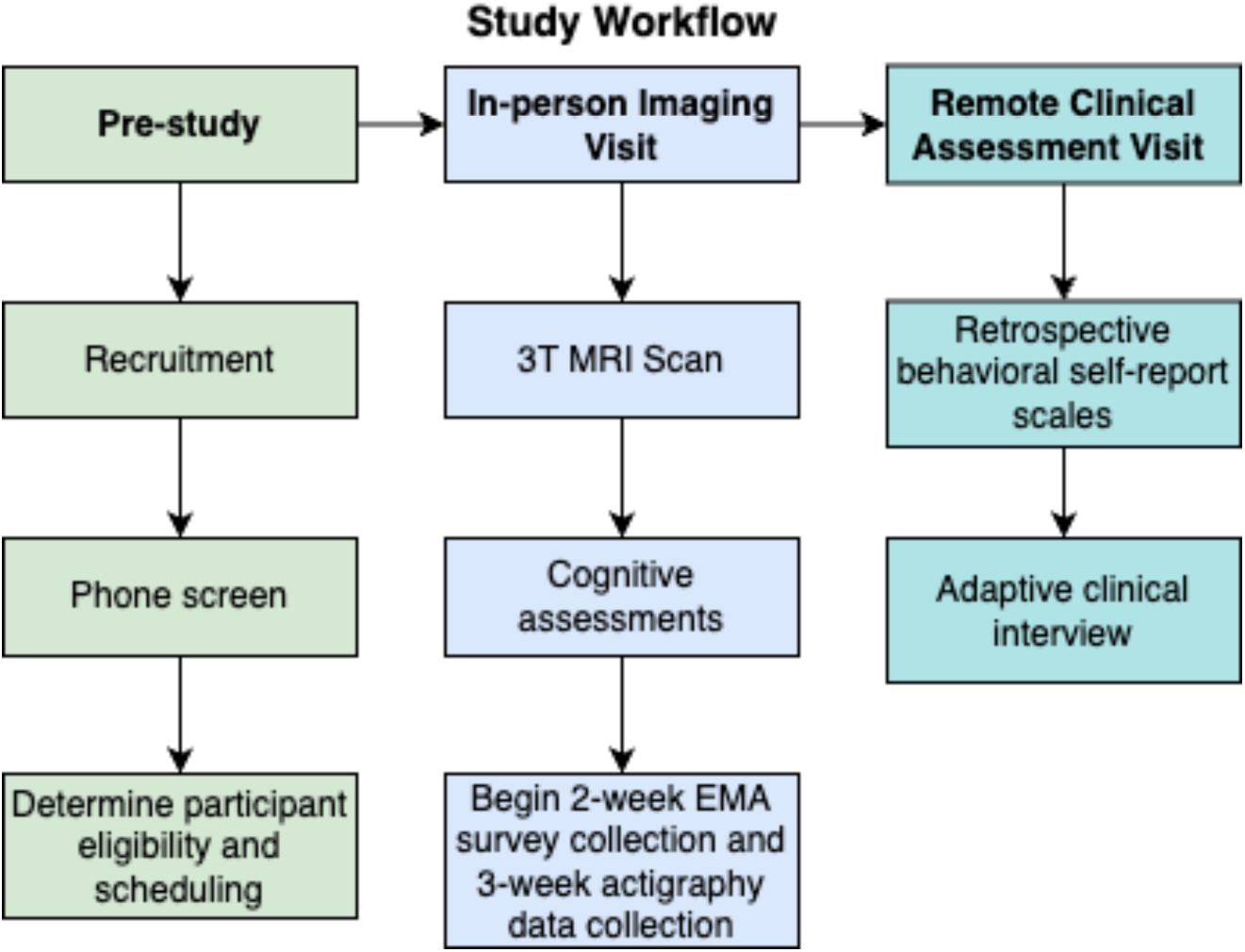
Study workflow.

### Self-Report Scales

A series of self-report scales assess demographic information, local environment, sleep, emotion regulation, and other domains of mental health. **Table 1** presents the self-report questionnaire battery administered during the remote clinical assessment visit.

**Table 1.**
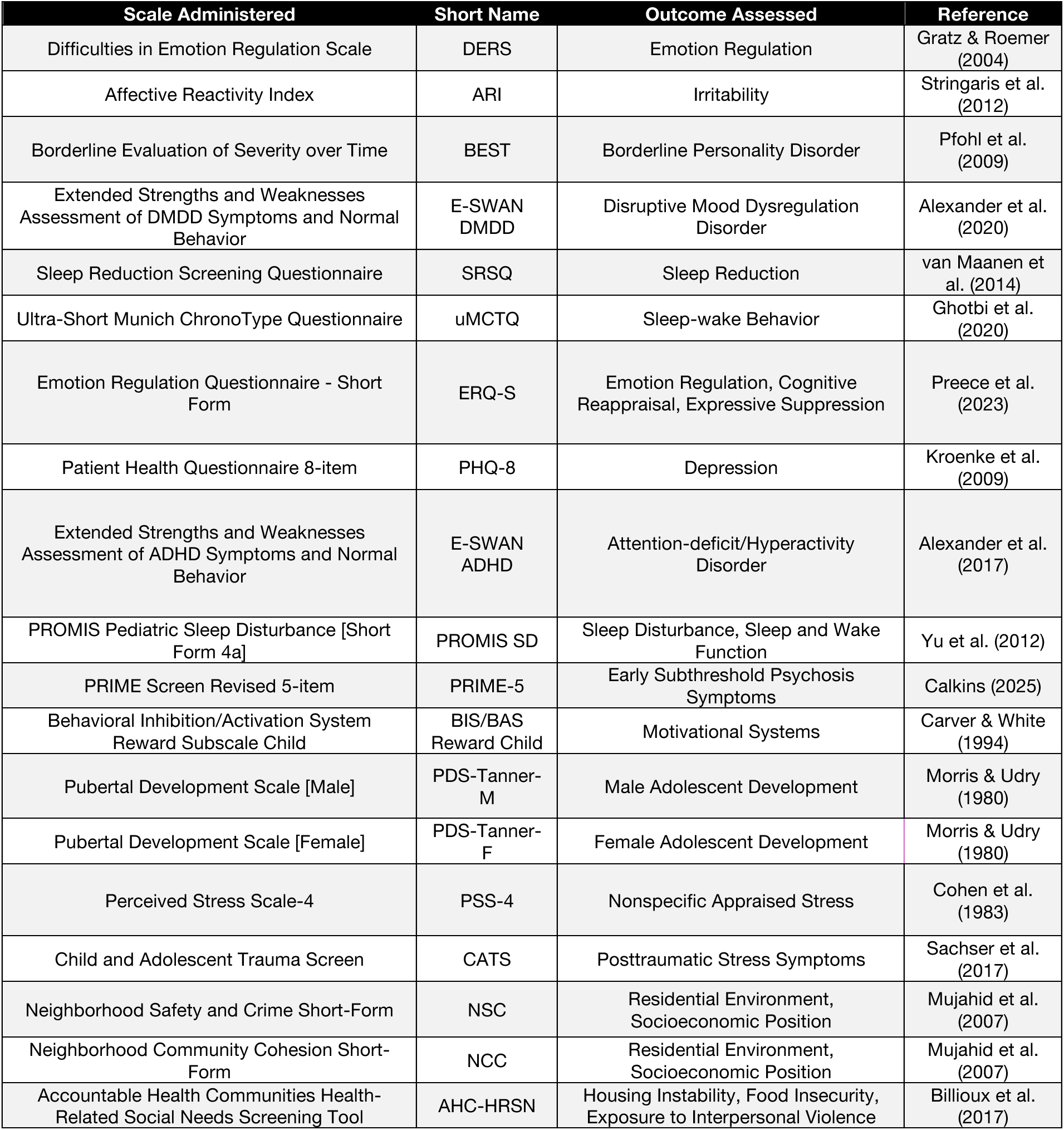
Self-report scales.

| Scale Administered | Short Name | Outcome Assessed | Reference |
| --- | --- | --- | --- |
| Difficulties in Emotion Regulation Scale | DERS | Emotion Regulation | Gratz & Roemer (2004) |
| Affective Reactivity Index | ARI | Irritability | Stringaris et al. (2012) |
| Borderline Evaluation of Severity over Time | BEST | Borderline Personality Disorder | Pfohl et al. (2009) |
| Extended Strengths and Weaknesses Assessment of DMDD Symptoms and Normal Behavior | E-SWAN DMDD | Disruptive Mood Dysregulation Disorder | Alexander et al. (2020) |
| Sleep Reduction Screening Questionnaire | SRSQ | Sleep Reduction | van Maanen et al. (2014) |
| Ultra-Short Munich ChronoType Questionnaire | uMCTQ | Sleep-wake Behavior | Ghotbi et al. (2020) |
| Emotion Regulation Questionnaire - Short Form | ERQ-S | Emotion Regulation, Cognitive Reappraisal, Expressive Suppression | Preece et al. (2023) |
| Patient Health Questionnaire 8-item | PHQ-8 | Depression | Kroenke et al. (2009) |
| Extended Strengths and Weaknesses Assessment of ADHD Symptoms and Normal Behavior | E-SWAN ADHD | Attention-deficit/Hyperactivity Disorder | Alexander et al. (2017) |
| PROMIS Pediatric Sleep Disturbance [Short Form 4a] | PROMIS SD | Sleep Disturbance, Sleep and Wake Function | Yu et al. (2012) |
| PRIME Screen Revised 5-item | PRIME-5 | Early Subthreshold Psychosis Symptoms | Calkins (2025) |
| Behavioral Inhibition/Activation System Reward Subscale Child | BIS/BAS Reward Child | Motivational Systems | Carver & White (1994) |
| Pubertal Development Scale [Male] | PDS-Tanner-M | Male Adolescent Development | Morris & Udry (1980) |
| Pubertal Development Scale [Female] | PDS-Tanner-F | Female Adolescent Development | Morris & Udry (1980) |
| Perceived Stress Scale-4 | PSS-4 | Nonspecific Appraised Stress | Cohen et al. (1983) |
| Child and Adolescent Trauma Screen | CATS | Posttraumatic Stress Symptoms | Sachser et al. (2017) |
| Neighborhood Safety and Crime Short-Form | NSC | Residential Environment, Socioeconomic Position | Mujahid et al. (2007) |
| Neighborhood Community Cohesion Short-Form | NCC | Residential Environment, Socioeconomic Position | Mujahid et al. (2007) |
| Accountable Health Communities Health-Related Social Needs Screening Tool | AHC-HRSN | Housing Instability, Food Insecurity, Exposure to Interpersonal Violence | Billieux et al. (2017) |

#### Pubertal measures

Participants under the age of 18 complete the Pubertal Developmental Scale,^51^ a self-report measure that uses Tanner’s images^52^ to assess pubertal status in adolescent populations. Male and female participants receive distinct versions of the scale. Both versions are seven items and include two items where participants indicate how their development compares to Tanner’s images; the items differ between male and female versions asking either about male pubertal changes (e.g., facial hair growth, deepening voice, etc.) or female pubertal changes (e.g., menstruation).

#### Measures of the childhood environment

Five scales assess childhood environment. First, the Childhood and Adolescent Trauma Screen (CATS)^53^ is a self-report questionnaire that examines participants’ exposure to potentially traumatic events (PTEs) and post-traumatic stress symptoms. An abbreviated 9-item version includes a structured PTE checklist asking about traumatic events (e.g., natural disasters, accidents, etc.) reflected in the DSM-5 criteria for post-traumatic stress disorder (PTSD). Furthermore, participants complete the Neighborhood Safety and Crime Short-Form (NSC; 3-item questionnaire)^54^ and the Neighborhood Community Cohesion Short-Form (NCC; 5-item questionnaire),^54^ two brief self-report scales that address neighborhood socioeconomic position. Finally, the Accountable Health Communities Health-Related Social Needs Screening Tool (AHC-HRSN)^55^ is a 14-item self-report scale that screens for non-medical health-related needs in the following domains: housing instability, food insecurity, transportation difficulties, utility assistance needs, interpersonal safety, financial strain, family and community support, education, physical activity, and disabilities. An additional 12 items are asked to obtain demographic and socioeconomic information from participants related to income, education, and household factors.

#### Self-report sleep measures

Three scales assess sleep. First, the Sleep Reduction Screening Questionnaire (SRSQ)^56^ is a 9-item self-report questionnaire measuring individual sleep need through daytime symptoms and interrogating sleep problems in order to assess perceived symptoms of sleep reduction (e.g., insufficient/poor sleep). Second, the Ultra-Short Munich ChronoType Questionnaire (uMCTQ)^57^ is a 6-item self-report questionnaire designed to evaluate sleep-wake-behavior. Finally, the PROMIS Pediatric Sleep Disturbance Short Form (PROMIS SD Short Form 4a)^58^ is a 4-item self-report measure evaluating sleep and wake function.

#### Measures of emotion lability, reactivity, and regulation

All participants complete an extensive battery including five complementary measures related to emotion regulation. First, the current study uses an abbreviated 5-item version of the Difficulties in Emotion Regulation Scale (DERS)^8^ assessing clinically relevant difficulties in emotion regulation. Second, the Emotion Regulation Questionnaire-Short Form (ERQ-S)^59^ is a 6-item self-report scale assessing two distinct emotion regulation strategies: cognitive reappraisal and expression suppression, designed for time-pressured settings. Third, the Affective Reactivity Index (ARI)^60^ is a 7-item self-report questionnaire that evaluates irritability over the past six months, focusing on feelings and behaviors specifically relevant to irritability, as well as overall impairment due to irritability. Fourth, the Extended Strengths and Weaknesses Assessment of DMDD Symptoms and Normal Behavior (E-SWAN DMDD)^61^ is a 30-item self-report questionnaire that measures elements of disruptive mood dysregulation disorder (DMDD). Fifth, participants complete a 10-item version of the Borderline Evaluation of Severity over Time (BEST)^62^ scale which measures symptoms of borderline personality disorder (BPD), including the Thoughts and Feelings (e.g., changes in self-identity, feelings of emptiness, etc.) and Behaviors-Negative (e.g., self-injury, impulsive sexual behavior, etc.) subscales. Sixth, participants complete the 5-item Reward Responsiveness subscale of the Behavioral Inhibition System and Behavioral Activation System Scales (BIS/BAS).^63^

#### Additional clinical self-report measures

In addition to detailed measures of emotion regulation, the study collects data on stressful life events, the symptoms of major depressive disorder (MDD), attention-deficit hyperactivity disorder (ADHD), and subthreshold symptoms of psychosis in three assessments. First, the Perceived Stress Scale-4 (PSS-4)^64^ is a 4-item self-report measure that assesses the degree of appraised stress associated with life events over the last month. Second, the Patient Health Questionnaire (PHQ-8)^65^ is an 8-item self-report measure of depression. Derived from the PHQ-9,^66^ the PHQ-8 includes all equivalent items except for item 9, which addresses thoughts of death or self-harm. Of note, item 9 was not administered due to issues of monitoring self-harm virtually. Third, the Extended Strengths and Weaknesses Assessment of ADHD Symptoms and Normal Behavior (E-SWAN ADHD)^61^ is an 18-item self-report questionnaire that measures ADHD. Lastly, the PRIME Screen Revised 5-item (PRIME-5)^67^ is a brief self-report measure that serves as an age-normed subthreshold psychosis screening tool.

#### Structured assessment of clinical domains

Participants complete the Computerized Adaptive Testing (CAT) GOASSESS^68^ psychopathology screener. The CAT GOASSESS is an adapted version of the highly-structured GOASSESS screening interview,^69^ and includes five domains: mood/anxiety, phobias, externalizing, psychosis, and pathological personality characteristics. While the full GOASSESS takes an hour on average to administer, the CAT GOASSESS is designed for rapid administration – often requiring only minutes – and minimal proctoring.^68^ **Table 2** compares the interview sections captured within the full and CAT GOASSESS versions.

**Table 2.**
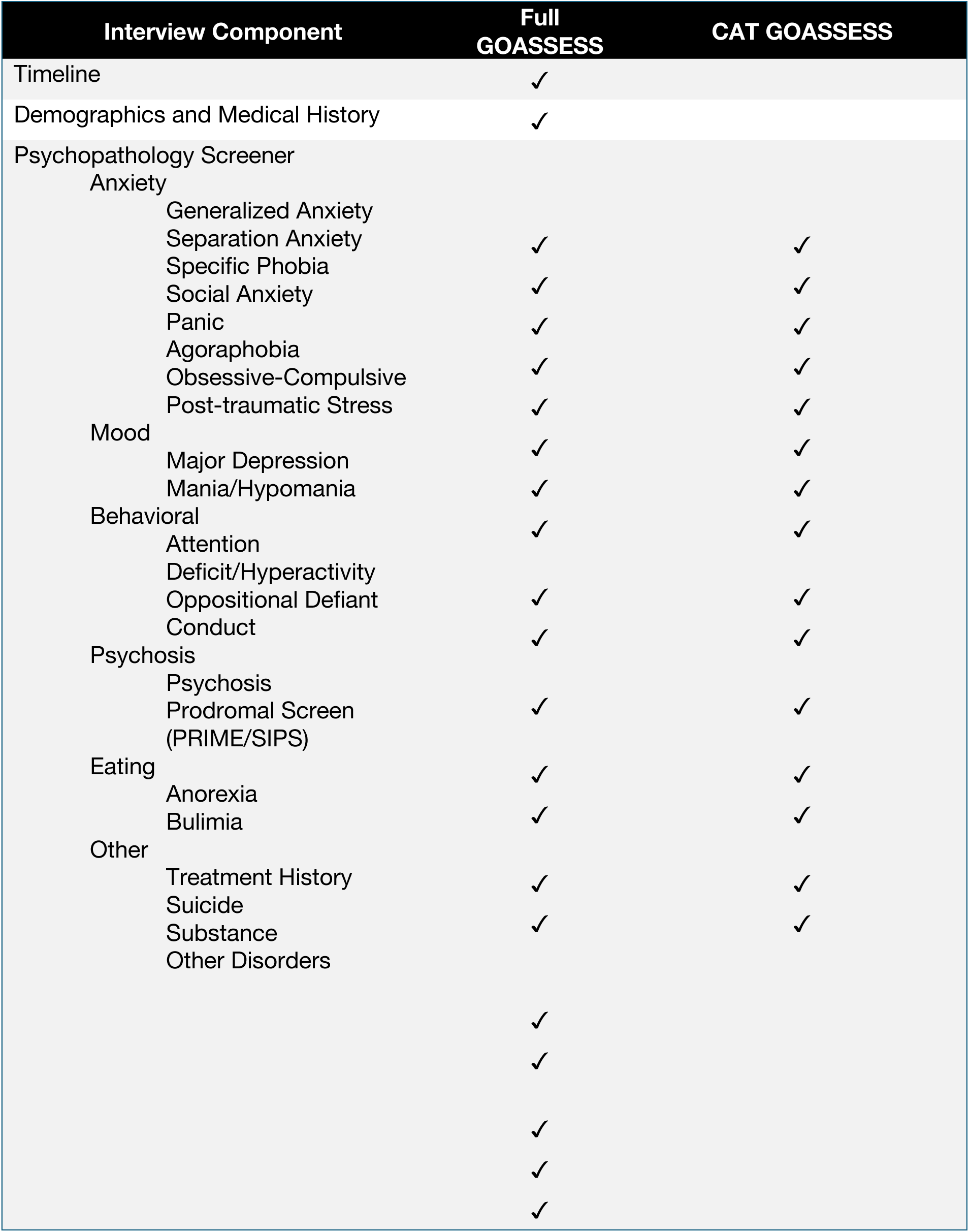
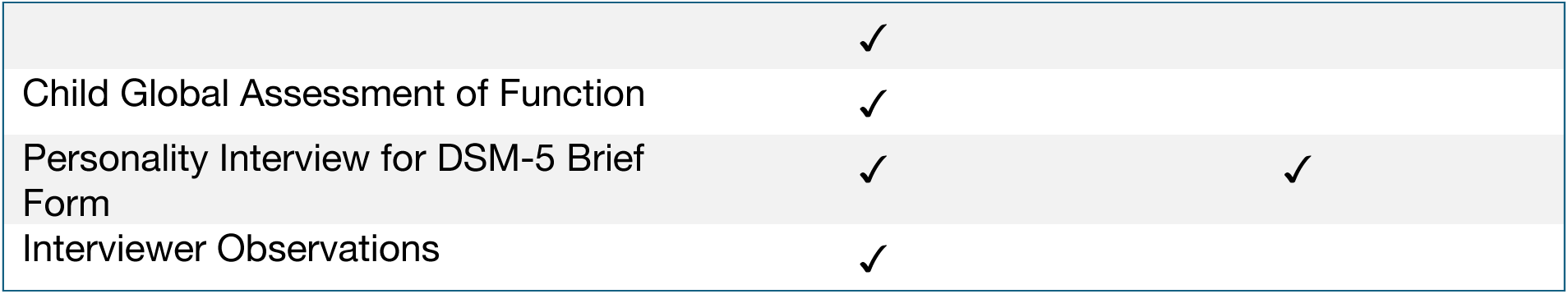
GOASSESS structured clinical interview. Elements of the Full and CAT versions of the GOASSESS are detailed.

### Cognitive Battery

We measure cognition using the Penn Computerized Neurocognitive Battery (CNB;^70^ see **Figure 2**). The CNB contains a series of computerized tests that are administered by a member of the research team. Participants typically complete the entire battery within an hour. The CNB evaluates performance accuracy and speed on neurobehavioral domains including: executive control, episodic memory, complex cognition, social cognition, sensorimotor speed, and reward decision-making.^71^ The CNB tests used in this study include the Abstraction, Inhibition, and Working Memory Test, Psychomotor Vigilance Test (3 Minute Version), Short Penn Continuous Performance Task (Adaptive), Digit Symbol Test, Penn Trailmaking Test (B Version), Short Letter-N-Back Test (2 Back Version), Short Visual Object Learning Test, Short Penn Logical Reasoning Test, Penn Emotion Recognition Task, and the Motor Praxis Test.

**Figure 2.**
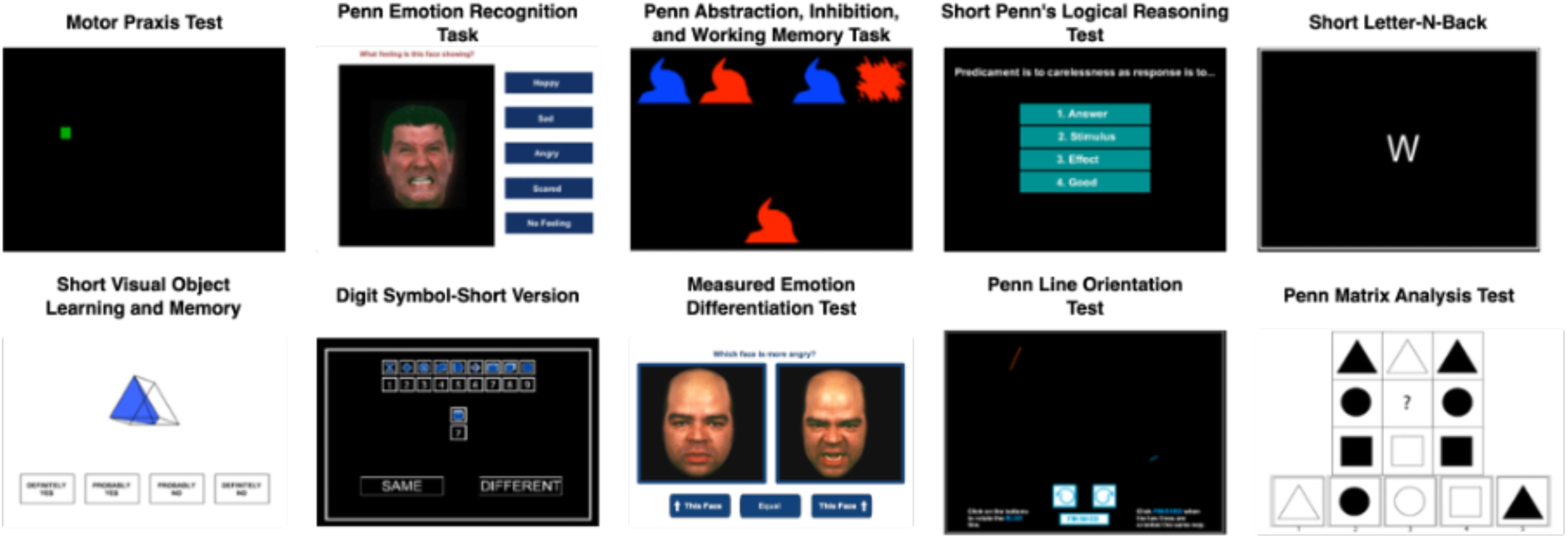
Cognitive battery. Exemplar CNB tasks featured across the CNB and CAT-CCNB batteries are displayed.

In addition to the CNB tasks described above, participants complete further cognitive assessments that use computerized adaptive testing (CAT) versions of the CNB. The CAT-CCNB evaluates the same domains, adding two measures of complex cognition (the Penn Matrix Analysis Test and the Penn Line Orientation Test), another measure of social cognition (the Measured Emotion Differentiation Test), and two reward/decision-making tests (Delay Discounting and Risk Discounting).^72^ See **Figure 2** for displays of ten exemplar cognitive tests included across the CNB and CAT-CCNB batteries; **Table 3** details the complete battery of cognitive tests and relevant domains.

**Table 3.**
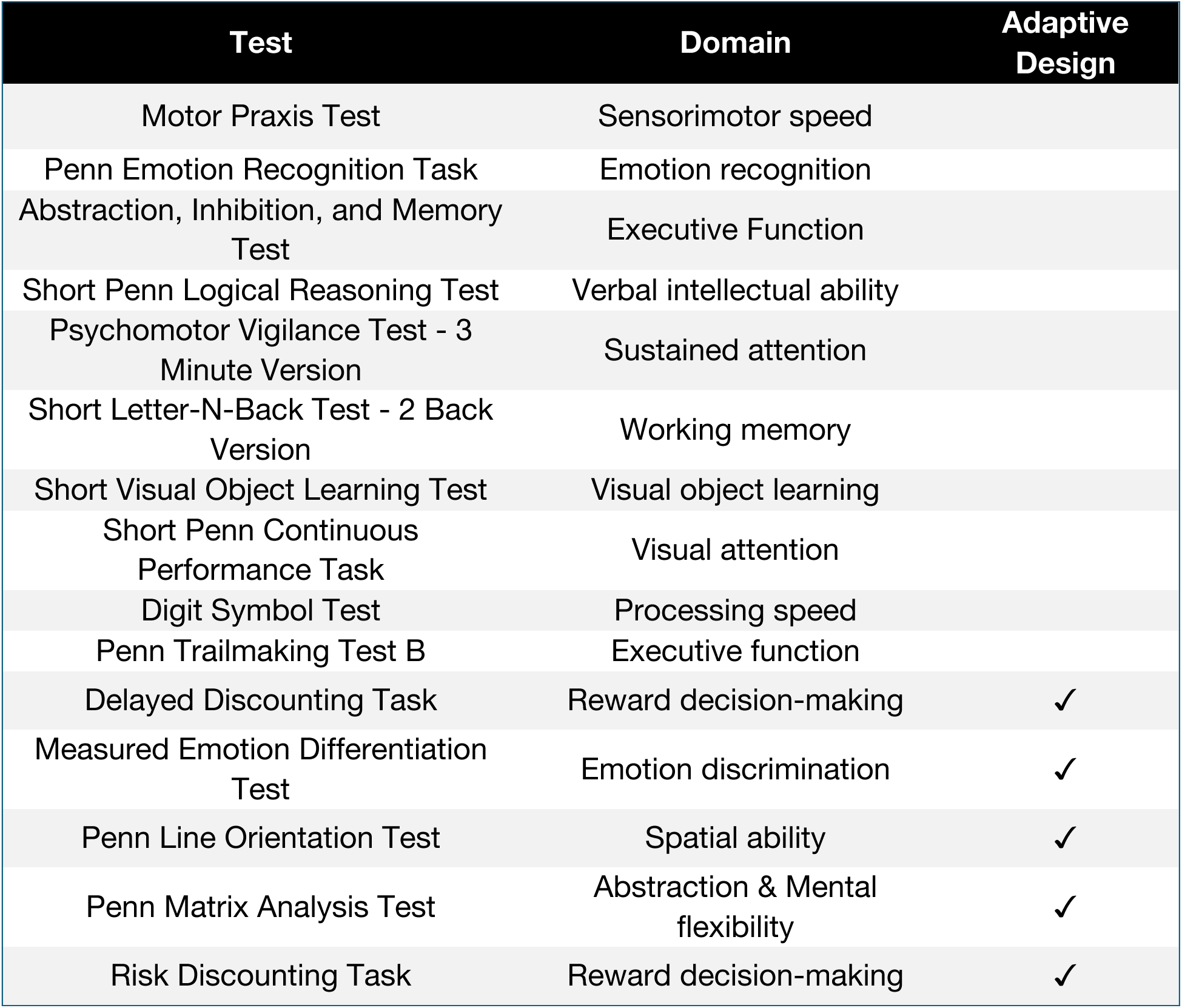
Cognitive battery domains.

### Ecological Momentary Assessment

Following the in-person imaging visit, participants begin a 2-week ecological momentary assessment (EMA) procedure with a mobile app entitled the “Real-time Ecological Assessment of the Context of mental and physical Health (*REACH*),” developed through a collaborative effort between the NIMH Intramural Research Program and the Child Mind Institute, adapted on the MindLogger Platform.^73^ MindLogger allows studies to design mental health assessments and interventions that are distributed to participants through customizable mobile or web activities.

EMA assesses participants’ emotions at the time of sampling rather than gathering a retrospective report on how they felt over the past week or month. Computerized EMA allows studies to verify the exact time when survey responses occurred;^74^ it thus reduces retrospective bias, provides real-time tracking of dynamic processes, and contextualizes relationships between symptoms and behaviors.^73^ *REACH* emerged after more than a decade of development and expansion of ecological assessment of emotional states with the mood circumplex^75^ as well as correlates of mood including sleep, physical activity, and energy.^76^ *REACH* includes modules on the mood circumplex, negative thoughts, sleep, context, screen and social media usage, life events, physical activity, food and drink intake, pain, and physical health. Descriptions of the specific items, procedures, and attribution are available through a Creative Commons License (Attribution-Non Commercial-Share Alike 4.0 International <u>Creative Commons license</u> agreement; CC BY-NC-SA 4.0). Versions of *REACH* are now being employed at several of the sites involved in an international collaborative effort on actigraphy and mood disorders (see below) that can enhance comparability and generalizability of the findings.

We configure the MindLogger app to send notification reminders prompting participants to complete the surveys at four daily time points: morning, mid-day, afternoon, and evening. Each participant determines the exact time when they receive these notification reminders based on their individual schedule. Participants have a 60-minute window to complete each survey. The MindLogger app allows participants to skip or exit the survey if they choose. See **Figure 3** for exemplar screenshots from MindLogger; the specific item prompts assigned across each of the four surveys are detailed in **Supplementary Table 1**.

**Figure 3.**
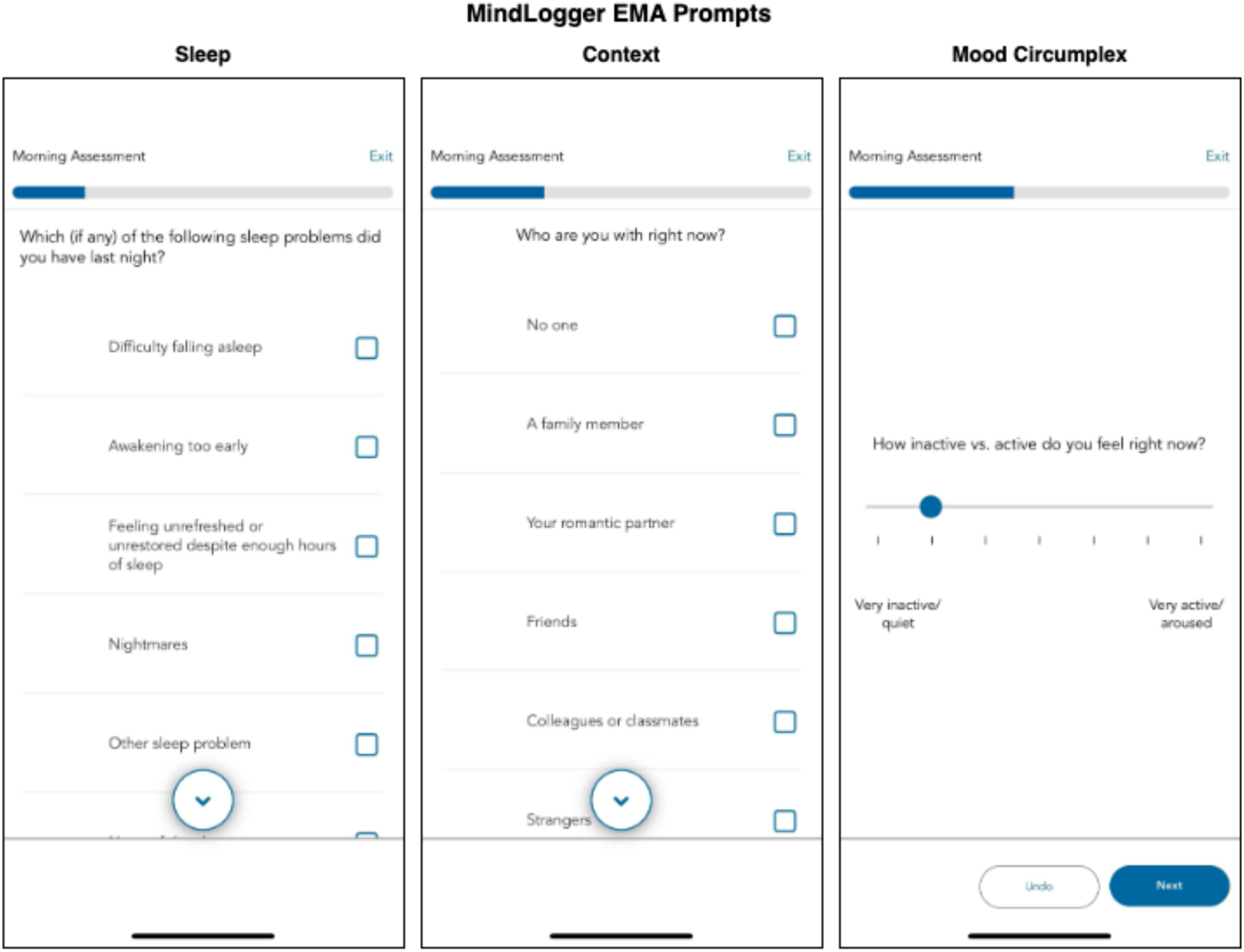
MindLogger EMA. Examples of three EMA prompts featured within the daily morning activity are displayed.

### Actigraphy

To measure physical activity and sleep,^77^ participants wear the GENEActiv actigraphy device (GENEActiv, Activinsights Ltd, Kimbolton, UK). The GENEActiv is a waterproof wristwatch-like device that participants are instructed to wear continuously for three weeks after the imaging visit. The GENEActiv devices are configured to collect raw acceleration data at 66.7 Hz for 21 days.

As part of our initial data release, we provide raw binary files containing acceleration data along the three x, y, and z movement axes. Binary files are extracted with the GENEActiv software version 4.0.12. Raw accelerometer data are processed with the GGIR R package (version 3.2.0),^78^ which performs auto-calibration, non-wear detection, and calculation of relevant sleep and physical activity variables. All processing is based on the Euclidean Norm Minus One (ENMO) metric with automated calibration. Acceleration angle metrics are calculated over 5-second epochs. Non-wear detection is performed using the 2023 algorithm, with an epoch length of 900 seconds for non-wear and signal clipping, and an epoch of 3600 seconds for non-wear detection. GGIR also evaluates the validity of each day and night of the study, requiring minimum valid wear time of 16 hours per day (midnight-midnight) and 12 hours per night (noon-noon). Sleep and wake are detected using the van Hees 2015 algorithm,^79,80^ which identifies sustained inactivity periods as intervals of at least five minutes during which the arm angle variability remains lower than five degrees. The sleep period window is detected using the HDCZA guider algorithm,^79^ which identifies the longest period of low movement and posture change to find the main daily sleep period. The sleep period time (SPT) is then used to calculate sleep variables, including sleep duration, defined as the total time spent in sustained inactivity bouts within the SPT window; wake after sleep onset (WASO), defined as the total time spent awake between the start and end of SPT; and sleep efficiency, calculated as the ratio of sleep duration to the total length of the SPT. Daytime physical activity levels are defined using GGIR’s default acceleration cutoff points of 40 mg for light, 100 mg for moderate, and 400 mg for vigorous activity. Sleep variables highlighted in the results section include total sleep duration (hours), WASO (hours), and sleep efficiency (%). Physical activity variables include time spent in light, moderate, and vigorous activity (in minutes). GGIR outputs several data quality variables, which we use to apply additional quality control. We first exclude recordings with device issues (e.g., clipping or failed calibration). Next, we exclude days or nights with poor data quality (e.g., >20% invalid data, >30% non-wear), as well as nights with abnormal features (extreme sleep durations < 2 or > 14 hours, extreme episode counts of < 5 or > 40, and daysleeper patterns). Finally, participants are excluded from each analysis if they have fewer than three valid days for activity analyses, and fewer than three valid nights for sleep analyses. Data from this study will be included in a collaborative effort with the Motor Activity Research Consortium for Health at the NIMH (mMARCH);^81^ that was established to harmonize the processing, data extraction, analytic methods and collection of common ancillary data for studies of mood disorders in adults and youth. In particular, mMARCH has made advances in processing accelerometry data^81^ and statistical analytic methods.^82^ Use of common procedures for actigraphy and EMA is now underway in several international settings (including the US, Brazil, Canada, Korea, and Switzerland) that will enhance the generalizability and interpretability of this research.

### Imaging Acquisition

MRI data is acquired on a 3T Siemens Magnetom Prisma (Erlangen, Germany) MRI scanner with the product 64-channel receiver array at the University of Pennsylvania. Sequence parameters and file naming conventions are summarized in **Table 4**; a full protocol PDF and the EXAR file is also available (see Data Availability, below).

**Table 4.** Imaging Parameters.

| Sequence | Slices | % FOV Phase | Resolution (mm) | TR (ms) | TE (ms) | TI (ms) | Flip Angle (°) | Multi Band Accel | Phase Partial Fourier |
| --- | --- | --- | --- | --- | --- | --- | --- | --- | --- |
| anat-T1w | 176 | 100% | 1.0x1.0x1.0 | 2500 | 2.9 | 1070 | 8 | N/A | Off |
| anat-T2w | 176 | 100% | 1.0x1.0x1.0 | 3200 | 565 | N/A | 120 | N/A | Allowed |
| fmap-epi_acq-func_dir-AP | 72 | 100% | 2.0x2.0x2.0 | 13690 | 20.80, 58.48, 96.16, 133.84, 171.52 | N/A | 90 | N/A | 6/8 |
| fmap-epi_acq-func_dir-PA | 72 | 100% | 2.0x2.0x2.0 | 13693 | 20.80, 58.48, 96.16, 133.84, 171.52 | N/A | 90 | N/A | 6/8 |
| func-bold_task-bao_dir-AP | 72 | 100% | 2.0x2.0x2.0 | 1761 | 14.20, 38.93, 63.66, 88.39, 113.12 | N/A | 68 | 6 | 6/8 |
| func-bold_task-rat_dir-PA | 72 | 100% | 2.0x2.0x2.0 | 1761 | 14.20, 38.93, 63.66, 88.39, 113.12 | N/A | 68 | 6 | 6/8 |
| dwi-dwi_acq-HASC92_dir-AP | 84 | 100% | 1.7x1.7x1.7 | 4300 | 90 | N/A | 90 | 4 | 7/8 |
| fmap-epi_acq-dwi_dir-PA | 84 | 100% | 1.7x1.7x1.7 | 4300 | 90 | N/A | 90 | 4 | 7/8 |
| perf-asl | 44 | 100% | 3.0x3.0x3.0 | 5000 | 10.8 | N/A | 120 | N/A | N/A |
| anat-MEGRE_acq-1p5mm | 96 | 75% | 1.5x1.5x1.5 | 35 | 7.5, 15, 22.5, 30 | N/A | 15 | N/A | Off |

#### Structural MRI

The imaging protocol begins with one run of T1-weighted MPRAGE structural MRI (named **anat-T1w**) that is aligned with the ABCD study^83^ (176 slices; repetition time, TR = 2500 ms; echo time, TE = 2.9 ms; flip angle, FA = 8°; field of view, FOV = 256×256 mm; matrix size = 256×256; voxel size = 1×1×1 mm; phase encoding direction = A >> P; acquisition time = 7:12). We also acquire the T2-weighted SPACE structural image (named **anat-T2w**) used in the ABCD protocol (176 slices; TR = 3200 ms; TE = 565 ms; FA = 120°; FOV = 256×256 mm; matrix size = 256×256; voxel size = 1×1×1 mm; phase encoding direction = A >> P; acquisition time 6:35). Both scans use embedded volumetric navigators to perform real-time motion correction, reducing motion artifacts in the resulting anatomical images.^84^

#### Functional MRI

This study includes two runs of multi-echo EPI. Previous work has shown that multi-echo EPI scans facilitate the identification of person-specific variation in BOLD fMRI with far less data.^48^ Before the fMRI time series are acquired, we collect PEpolar-style field maps for distortion correction of the fMRI data. These field maps (named **fmap-epi_acq-func_dir-AP** and **fmap-epi_acq-func_dir-PA**) consist of two multi-echo EPI acquisitions with the same acquisition parameters as the fMRI scans with opposite phase encoding directions (72 slices; TR = 13,690 ms; TE = 20.80, 58.48, 96.16, 133.84, 171.52 ms; FA = 90°; FOV = 220×220 mm; matrix size = 110×110; voxel size = 2×2×2 mm; acquisition time = 1:22). Current software cannot leverage multiple echoes in PEpolar-style field maps, so we retain the first echo from each multi-echo field map with **‘**acq-func**’** (e.g., **‘**sub-01_ses-01_acq-func_dir-AP_epi.nii.gz**’**). The multi-echo versions of the field maps are retained in the dataset for potential future use.

Two runs of functional MRI (named **func-bold_task-bao_dir-AP** and **func-bold_task-rat_dir-PA**) are acquired with a multiband EPI sequence (CMRR, University of Minnesota). Each functional run is acquired with the same protocol (72 slices; TR = 1761 ms; TE = 14.20, 38.93, 63.66, 88.39, 113.12 ms; FA = 68°; FOV = 220×220 mm; matrix size = 110×110; voxel size = 2×2×2 mm; multiband factor = 6; in-plane acceleration factor = 2). This protocol is highly similar to the one used by Moser et al. (2025)^85^ and Siegel et al. (2024).^86^ During each run, participants view Pixar short animated movies: “Bao” and “Your Friend the Rat.” For “Bao,” the phase encoding direction is anterior-to-posterior and each run is 7:13, during which 246 volumes are acquired. For “Your Friend the Rat,” the phase encoding direction is posterior-to-anterior and each run is 10:45 minutes long, during which 366 volumes are acquired. For both runs, a single-band reference image is also acquired for each echo to assist with co-registration. Furthermore, three no-excitation noise volumes are also acquired at the end of each run to allow for implementation of denoising with NORDIC.^85,87,88^ Finally, we acquire all fMRI data with both magnitude and phase reconstruction enabled; this complex data is helpful for denoising with NORDIC, improves T2* estimation, and allows for cutting edge distortion correction methods (e.g., MEDIC).^89^

#### Diffusion MRI

This study includes compressed-sensing diffusion spectrum imaging (CS-DSI). The vast majority of dMRI images sample many directions at one or more *b-*values (or shells). In contrast, DSI scans densely sample *q-*space on a Cartesian grid. This enables the direct estimation of the diffusion Ensemble Average Propagator (EAP), the physical process driving biologically meaningful derivatives of dMRI.^90^ Previous research has demonstrated that DSI achieves greater biological fidelity in tractography when compared to ground-truth anatomic tracings through more accurate ODF estimates.^91,92^ DSI scans have also been shown to result in improved tractography^93^ and enhanced gray-white contrast.^94^ While available research emphasizes massive potential for DSI, the dense Cartesian sampling requires very long scan times (30+ minutes), which has prohibited the wide use of DSI.

To mitigate the need for long DSI scan times, recent work has shown that CS-DSI acquisitions can undersample *q*-space and still accurately estimate the underlying EAPs.^95–100^ Our team recently published the first in-depth study that compared CS-DSI and full DSI scans in humans.^50^ We found that a 7.4-minute CS-DSI scan provided highly similar accuracy and reliability to a full DSI scan. This study acquires a 92-direction CS-DSI scan (named **dwi-dwi_acq-HASC92_dir-AP**) that uses a homogenous angular sampling scheme (HA-SC;^101^ TR = 4300 ms; TE = 90.00 ms; FOV = 230×230 mm; voxel size = 1.7×1.7×1.7 mm; 84 interleaved slices acquired anterior to posterior; a multiband factor of 4; acquisition time = 9:02).^98^ In addition to 95 diffusion-weighted images, 9 *b* = 0 images are acquired (104 volumes total). For full details on the CS-DSI sampling schemes see Data Availability, below. dMRI data is acquired with both magnitude and phase reconstruction enabled; this complex data is helpful for denoising with Marchenko-Pastur principal components analysis (MP-PCA) (as described below). For distortion correction, we acquire a reverse phase-encoded scan (named **fmap-epi_acq-dwi_dir-PA**) that includes 7 volumes (TR = 4300 ms; TE = 90.00 ms; FOV = 230×230 mm; voxel size = 1.7×1.7×1.7 mm; 84 interleaved slices acquired posterior to anterior; multiband factor = 4; acquisition time = 2:09).

#### Arterial spin-labeled MRI

To measure brain perfusion, we acquire a state-of-the-art arterial spin-labeled (ASL) MRI scan. Specifically, we use a background-suppressed unbalanced PCASL scan (named **perf-asl**) using a stack of spirals turbo spin echo (Labeling duration = 1.8 s, Post-labeling delay (PLD) = 1.8s, TR = 5000.0 ms; TE = 10.8 ms; FOV = 240×240 mm; voxel size = 3.0×3.0×3.0 mm; acquisition time = 4:24).^102,103^ We also acquire a reference scan to assist in ASL calibration (M0 and T1 estimation) and co-registration, using the same sequence to acquire a presaturated, unsuppressed proton-density weighted volume (Tsat = 5 s) and presaturated inversion recovery volume (Tsat = 5 s, TI = 1.978 s). This reference scan can be used to calculate a low-resolution quantitative T1 map.^104^

#### MEGRE

We acquire one run of multi-echo gradient-recalled echo (MEGRE; named **anat-MEGRE_acq-1p5mm**) scan with magnitude and phase reconstruction (96 slices; repetition time, TR = 35 ms; echo times, TEs = 7.5, 15, 22.5, 30 ms; flip angle, FA = 15°; field of view, FOV = 180×240 mm; matrix size = 120×160; voxel size = 1.5×1.5×1.5 mm; phase encoding direction=A >> P; acquisition time = 3:59). This scan can be used for quantitative susceptibility mapping (QSM), which is sensitive to developmental changes in brain iron and myelin.

### Image Processing

As part of our initial data release, we provide processed T1-weighted, fMRI, dMRI, and ASL images. sMRIPrep is used for processing T1-weighted (T1w) and T2-weighted (T2w) images. fMRIPrep is used to minimally preprocess fMRI data;^105^ fMRI post-processing is performed with fMRIPost-AROMA,^106^ tedana,^107^ and XCP-D.^108^ dMRI is processed with QSIPrep and QSIRecon.^109^ ASL images are processed with ASLPrep.^110^ As of the current data release, we do not provide processed MEGRE/QSM images.

#### Structural image processing

Preprocessing of T1-weighted images uses sMRIPrep 0.16.0, as implemented in fMRIPrep 24.1.1^105^ using Nipype 1.8.6.^111,112^ The T1w image undergoes correction for intensity non-uniformity with N4BiasFieldCorrection with ANTs 2.5.3,^113,114^ skull-stripping with a Nipype implementation of the ANTs brain extraction workflow, and brain tissue segmentation with FSL’s FAST 6.0.7.7.^115^ Brain surfaces are reconstructed using FreeSurfer 7.3.2,^116^ and the brain mask estimated previously is refined with a custom variation of the method to reconcile ANTs-derived and FreeSurfer-derived segmentations.^117^ The T2-weighted image is used to improve pial surface refinement. Volume-based spatial normalization to two standard spaces (MNI152NLin6Asym and MNI152NLin2009cAsym, accessed through TemplateFlow 24.2.0)^118^ is performed through nonlinear registration with antsRegistration (ANTs 2.5.3), using brain-extracted versions of both T1w reference and the T1w template. A CIFTI grayordinate file containing 91k vertices is resampled onto the fsLR template using Connectome Workbench.^119,120^

#### Functional image preprocessing

fMRI data is preprocessed using fMRIPrep 24.1.1,^105,121^ which is based on Nipype 1.8.6.^111,112^ First, a *B_0_*-nonuniformity map is estimated based on echo-planar imaging (EPI) references using FSL’s TOPUP.^122^ For each BOLD run, the following preprocessing is performed. First, a reference volume is generated from the shortest echo of the BOLD run for use in head-motion correction. Head-motion parameters with respect to the BOLD reference (transformation matrices, and six corresponding rotation and translation parameters) are estimated before any spatiotemporal filtering using FSL’s mcflirt.^123^ The estimated fieldmap is aligned with rigid-registration to the target EPI reference run. The BOLD reference is then co-registered to the T1w reference using FreeSurfer’s bbregister^124^ with six degrees of freedom. The aligned T2w image is also used for initial co-registration. The BOLD time series are resampled onto the left/right-symmetric template “fsLR” using Connectome Workbench.^119,120^ A “goodvoxels” mask is applied during volume-to-surface sampling in fsLR space, excluding voxels whose time series have a locally high coefficient of variation. Grayordinates files^119^ containing 91k samples are also generated with surface data transformed directly to fsLR space and subcortical data transformed to 2 mm resolution MNI152NLin6Asym space.

#### Post-processing of functional images

Following preprocessing with fMRIPrep, images are post-processed with complementary software. We seek to decompose the processed fMRI time series into “noise” and “signal” components using two methods. First, ICA-AROMA^106,125^ is implemented using fMRIPost-AROMA 0.0.10, which is based on Nipype 1.9.2.^111,112^ Noise and signal components from ICA-AROMA are compared to tedana,^126^ which is run using the “minimal” decision tree – a simplified version of the MEICA decision tree. Tedana fits a monoexponential model to the data at each voxel using nonlinear model fitting to estimate T2* and S0 maps, using T2*/S0 estimates from a log-linear fit as initial values. For each voxel, the value from the adaptive mask is used to determine which echoes would be used to estimate T2* and S0. In cases of model fit failure, T2*/S0 estimates from the log-linear fit are retained instead. Multi-echo data are then optimally combined using the T2* combination method.^127^ Next, component selection is performed to identify BOLD (TE-dependent) and non-BOLD (TE-independent) components using a decision tree. Rejected components’ time series are then orthogonalized with respect to accepted components’ time series. Minimum image regression is then applied to the data to remove spatially diffuse noise.^126^ The tedana workflow uses NumPy,^128^ SciPy,^129^ pandas,^130^ scikit-learn,^131^ Nilearn,^132^ Bokeh,^133^ Matplotlib,^134^ and Nibabel.^135^

After tedana is applied, the component classifications from fMRIPost-AROMA and tedana are combined, such that any component that is rejected by either ICA-AROMA or ME-ICA is labeled as rejected. The rejected components’ time series are then orthogonalized with respect to accepted components’ time series. Minimum image regression is then applied to the mixing matrix in order to remove spatially diffuse noise.^126^

Following identification of noise components using ICA-AROMA and tedana, images are processed using XCP–D.^108,136,137^ XCP-D is built with Nipype 1.9.2.^111,112^ Many internal operations of XCP-D use AFNI,^138,139^ Connectome Workbench,^119,120^ ANTS,^140^ TemplateFlow 24.2.2,^118^ Matplotlib 3.10.0,^134^ Nibabel 5.3.2,^135^ Nilearn 0.11,^132^ NumPy 2.2.1,^141^ pybids 0.18.1,^142^ and SciPy 1.15.1.^129^

Non-steady-state volumes are extracted from the preprocessed confounds and are discarded from both the BOLD data and nuisance regressors. The set of orthogonalized nuisance components from the tedana step is selected for the nuisance regression, along with mean white matter signal, mean cerebrospinal fluid signal, and mean global signal.^137,143^ The BOLD data are converted to NIfTI format, despiked with AFNI’s 3dDespike, and converted back to CIFTI format. Nuisance regressors are regressed from the BOLD data using a denoising method based on Nilearn’s approach. The time series are band-pass filtered using a second-order Butterworth filter, in order to retain signals between 0.01-0.08 Hz. The same filter is applied to the confounds. The resulting time series are then denoised using linear regression. The denoised BOLD is then smoothed using Connectome Workbench with a Gaussian kernel (FWHM = 6 mm).

Processed functional time series are extracted from residual BOLD using Connectome Workbench^119,120^ for the atlases. Corresponding pairwise functional connectivity between all regions is computed for each atlas, which is operationalized as the Pearson’s correlation of each parcel’s unsmoothed time series with the Connectome Workbench. In cases of partial coverage, uncovered vertices (values of all zeros or NaNs) are either ignored (when the parcel had >50.0% coverage) or the whole parcel is set to zero (when the parcel had <50.0% coverage). The following atlases are used in the workflow: the Schaefer Supplemented with Subcortical Structures (4S) atlas^119,144–147^ at 10 different resolutions (156, 256, 356, 456, 556, 656, 756, 856, 956, and 1056 parcels), the Glasser atlas,^148^ the Gordon atlas,^149^ the Tian subcortical atlas,^150^ the HCP CIFTI subcortical atlas,^119^ and the MIDB precision brain atlas derived from ABCD data and thresholded at 75% probability.^151^ In addition, XCP-D calculates two scalar maps. First, the amplitude of low-frequency fluctuation (ALFF)^152^ is computed by transforming the mean-centered, standard deviation-normalized, denoised BOLD time series to the frequency domain.

The power spectrum is computed within the 0.01-0.08 Hz frequency band and the mean square root of the power spectrum is calculated at each voxel to yield voxel-wise ALFF measures. The resulting ALFF values are then multiplied by the standard deviation of the denoised BOLD time series to retain the original scaling. The ALFF maps are smoothed with the Connectome Workbench using a Gaussian kernel (FWHM = 6 mm). Second, for each hemisphere, regional homogeneity (ReHo)^153^ is computed using surface-based 2dReHo.^154^ Specifically, for each vertex on the surface, Kendall’s coefficient of concordance (KCC) is computed with nearest-neighbor vertices to yield ReHo. For the subcortical, volumetric data, ReHo is computed with neighborhood voxels using AFNI’s 3dReHo.^155^

#### dMRI Processing

dMRI preprocessing is performed using *QSIPrep* 1.0.0,^109^ which is based on Nipype 1.9.1;^111,112^ (RRID:SCR_002502). Many internal operations of QSIPrep use Nilearn 0.10.1^132^ (RRID:SCR_001362) and Dipy.^156^ QSIPrep uses a slightly different anatomical workflow than sMRIPrep/fMRIPrep. Specifically, the T1w image is corrected for intensity non-uniformity (INU) and used as the T1w-reference map. The anatomical reference image is first reoriented into AC-PC alignment using the 6-degrees-of-freedom (rigid) component of a full affine registration to MNI152NLin2009cAsym, and a symmetric nonlinear registration (SyN) using antsRegistration to further refine alignment to the MNI152NLin2009cAsym template. Brain extraction is performed on the T1w image using SynthStrip^157^ and automated segmentation is performed using SynthSeg^158^ from FreeSurfer version 7.3.1.

Next, any diffusion images with a b-value less than 100 s/mm^2^ are treated as a *b* = 0 image. Magnitude and phase diffusion-weighted imaging data are combined into a complex-valued file, then denoised using the Marchenko-Pastur PCA method implemented in MRtrix3’s dwidenoise^159–161^ with a 3-voxel window. After denoising, the complex-valued data are split back into magnitude and phase. The mean intensity of the diffusion-weighted series is adjusted so all the mean intensity of the *b* = 0 images matched across each separate DWI scanning sequence.

Initial motion correction is performed using only the *b* = 0 images. An unbiased *b* = 0 template is constructed over two iterations of 3dSHORE registrations. To estimate head motion in b>0 images, the SHORELine method^50,109^ is used to iteratively leave out each b>0 image and reconstruct the remaining images using 3dSHORE;^96^ the signal for the left-out image served as the registration target. The reconstructed, model-generated images are transformed into alignment with each b>0 image. Both slicewise and whole-brain quality control measures (cross correlation and *R*^2^) are calculated.

Susceptibility distortion correction is performed using DRBUDDI,^162^ part of the TORTOISE^163^ software package. Reverse phase-encoding EPI-based fieldmaps are collected, resulting in pairs of images with distortions going in opposite directions. DRBUDDI uses *b* = 0 reference images with reversed phase encoding directions to estimate the susceptibility-induced off-resonance field. A T2-weighted image is included in the multimodal registration. The DWI time series are resampled to AC-PC, generating a preprocessed DWI run in AC-PC space with 1.7 mm isotropic voxels.

Following preprocessing, the preprocessed dMRI images are reconstructed using QSIRecon (1.0.1).^109^ Diffusion orientation distribution functions (ODFs) are reconstructed using generalized q-sampling imaging (GQI)^164^ with a ratio of mean diffusion distance of 1.250000 in DSI Studio (version 94b9c79). Automatic Tractography is run in DSI Studio (version 94b9c79) and bundle shape statistics are calculated.^165^

#### ASL image processing

Arterial spin-labeled MRI images are preprocessed using ASLPrep 0.7.5,^110,166^ which is based on fMRIPrep^105,167^ and Nipype 1.8.6.^111^ Many internal operations of ASLPrep use Nilearn 0.11.1,^132^ NumPy,^141^ and SciPy.^129^ In large part, ASLPrep uses the precomputed structural image processing from sMRIPrep, as described above. The ASL reference scan is co-registered to the T1w reference using Freesurfer’s bbregister which implements boundary-based registration^124^ with six degrees of freedom. All resampling in ASLPrep uses a single interpolation step that concatenates all transformations. Gridded (volumetric) resampling is performed using antsApplyTransforms, configured with Lanczos interpolation to minimize the smoothing effects of other kernels.^168^

Head-motion parameters are estimated for the ASL data using FSL’s mcflirt.^123^ Motion correction is performed separately for label and control volumes in order to account for intensity differences between different contrasts; when these volumes are motion corrected together, intensity differences can be conflated with head motions.^169^ Next, ASLPrep concatenates the motion parameters across volume types and re-calculates relative root mean-squared deviation. Several confounding time series are calculated, including both framewise displacement (FD) and DVARS. FD and DVARS are calculated using the implementations in Nipype (following the definition by Power et al., 2014^170^) for each ASL run. ASLPrep summarizes in-scanner motion as the mean framewise displacement and relative root-mean square displacement.

ASLPrep is used to calculate cerebral blood flow (CBF) from the single-delay PCASL using a single-compartment general kinetic model.^171^ Calibration (M0) volumes associated with the ASL scan are smoothed with a Gaussian kernel (FWHM = 5 mm) and the average calibration image is calculated and scaled by 10.0. The quality evaluation index (QEI) is computed for each CBF map.^172^ QEI is based on the similarity between the CBF and the structural images, the spatial variability of the CBF image, and the percentage of gray matter voxels containing negative CBF values.

Parcellated CBF estimates are extracted for multiple atlases, including the Schaefer Supplemented with Subcortical Structures (4S) atlas^119,144–147^ at 10 different resolutions (156, 256, 356, 456, 556, 656, 756, 856, 956, and 1056 parcels), the Glasser atlas,^148^ the Gordon atlas,^149^ the Tian subcortical atlas,^150^ and the HCP CIFTI subcortical atlas.^119^ In cases of partial coverage, either uncovered voxels (values of all zeros or NaNs) are ignored (when the parcel has >50.0% coverage) or the whole parcel is set to zero (when the parcel has <50.0% coverage).

## Results

Here we present initial data from the first participants enrolled in the study protocol. As described below, we focus on EMA, actigraphy, and neuroimaging data. Notably, all processed data described below are openly available on OpenNeuro; all processing code and software is available on GitHub (see Data and Code Availability, below). Data will be updated regularly as study accrual continues.

### Ecological Momentary Assessment (EMA)

A total of n = 7 completed the 14-day EMA procedure in this preliminary data release; EMA was not available for the first three study participants enrolled. We examined participant compliance with the EMA procedure according to time of day over the study period by calculating the percentage of prompts to which participants responded for each time point (**Figure 4**). The current study’s EMA adherence averaged 72% across all activities, with compliance of 74% in the morning, 82% at mid-day, 82% in the afternoon, and 71% in the evening. In addition to adherence, we examined mean scores and daily patterns in mood circumplex items (**Figures 5 & 6**). Out of a 7-point scale, we found a mean sadness score of 3.50 (SD = 0.91), a mean anxiousness score of 3.67 (SD = 0.96), a mean positive thoughts score of 4.30 (SD = 1), a mean negative thoughts score of 3.49 (SD = 0.86), a mean active score of 3.20 (SD = 1.16), and a mean tiredness score of 3.46 (SD = 1.18).

**Figure 4.**
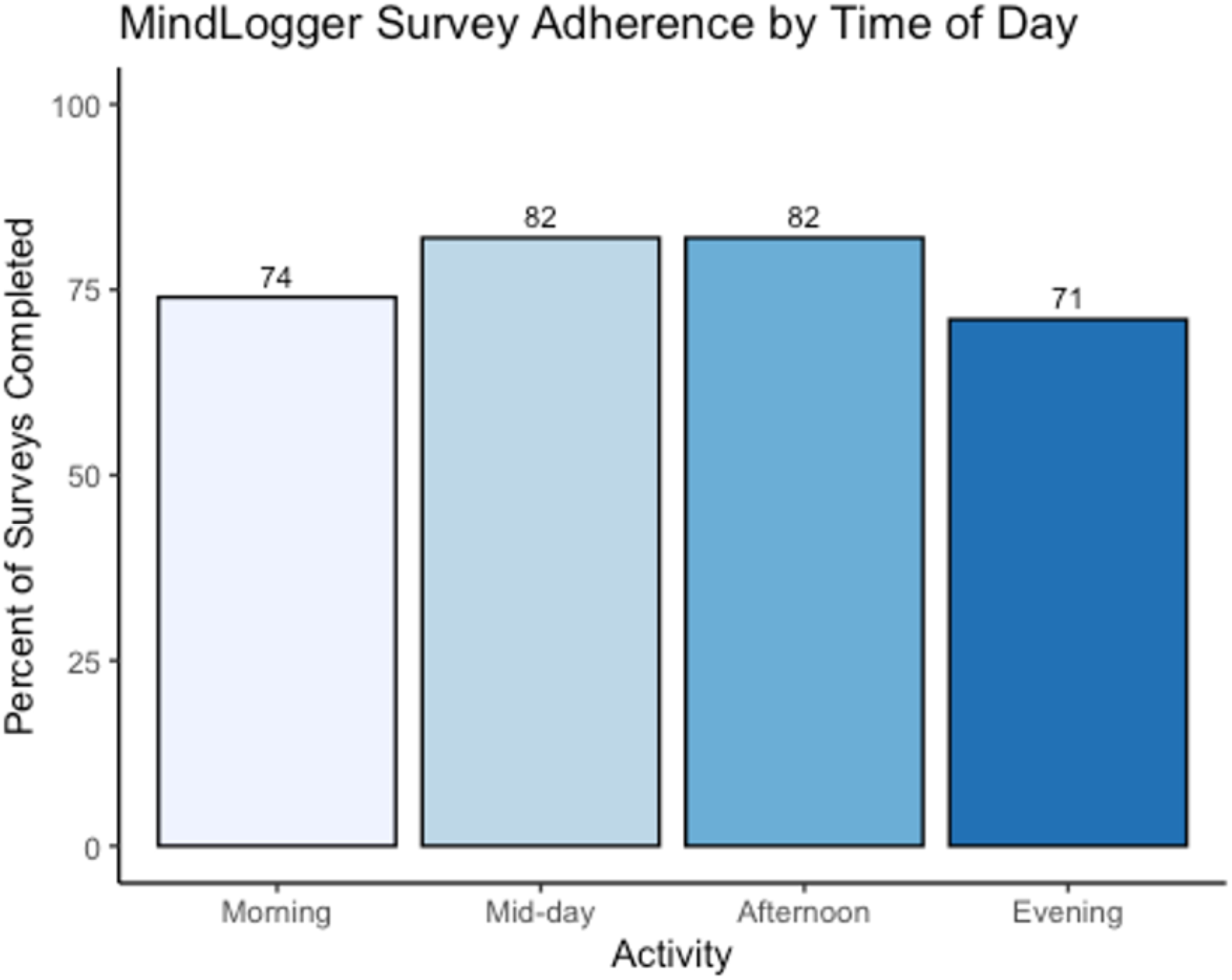
Ecological momentary assessment (EMA) adherence. Average adherence for each survey across the 14-day EMA period is displayed for each of the four daily timepoints.

**Figure 5.**
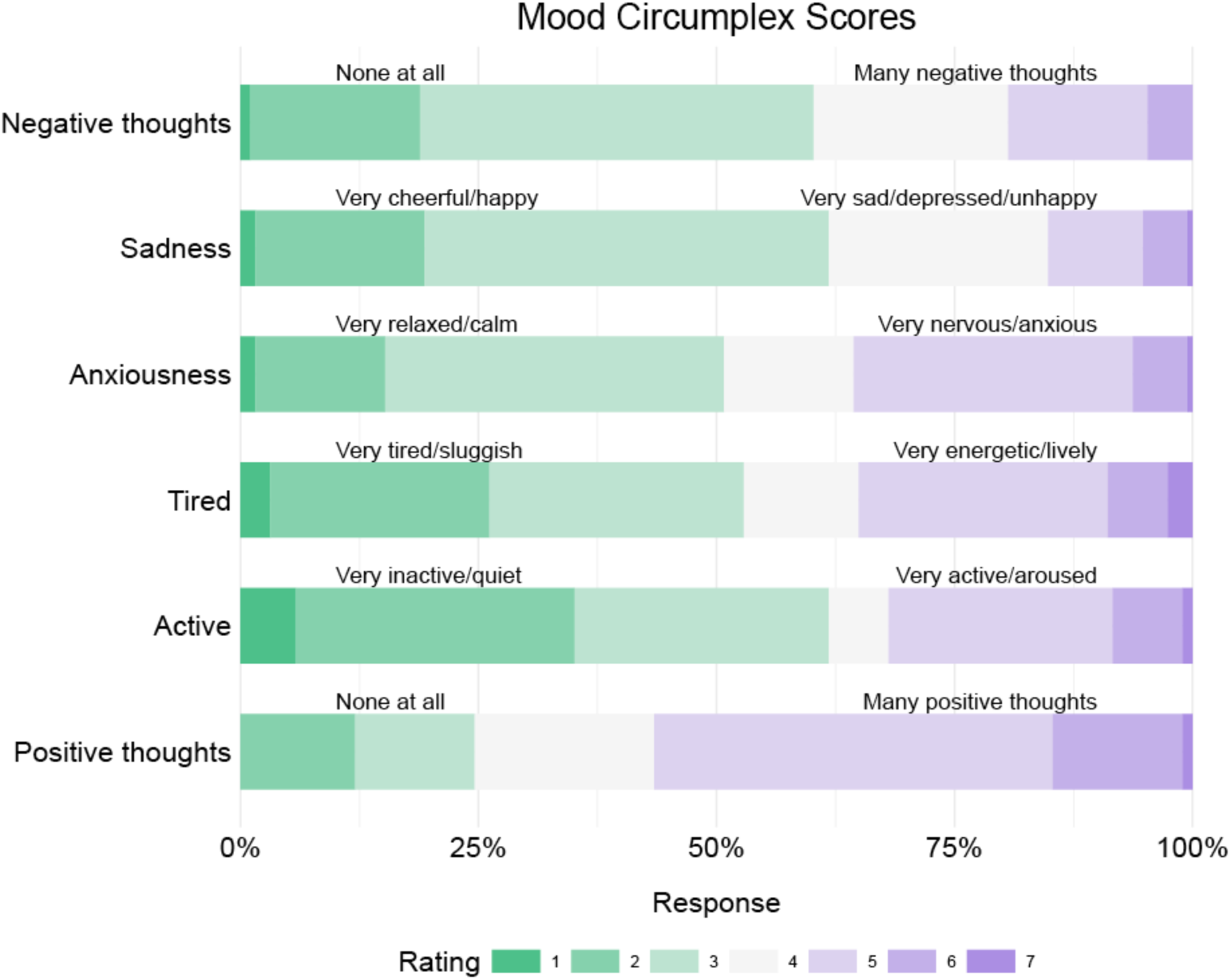
Ecological momentary assessment (EMA) mood scores. The proportion of mood circumplex scores across participants and timepoints are displayed. Each mood score is represented by a horizontal bar, with colors indicating rating intensity on a 1-7 scale. The length of each colored segment shows the proportion of responses at each rating level across participants and activity types (morning, mid-day, afternoon, and evening assessments). Green represents lower ratings (1-2), light gray represents neutral ratings (3-5), and purple represents higher ratings (6-7). The specific meaning of high and low ratings varies for each measure as indicated by the labels on either side of the bars. For example, in ‘Sadness,’ a longer green segment indicates more responses of ‘Very cheerful/happy’ while a longer purple segment indicates more responses of ‘Very sad/depressed/unhappy.’

**Figure 6.**
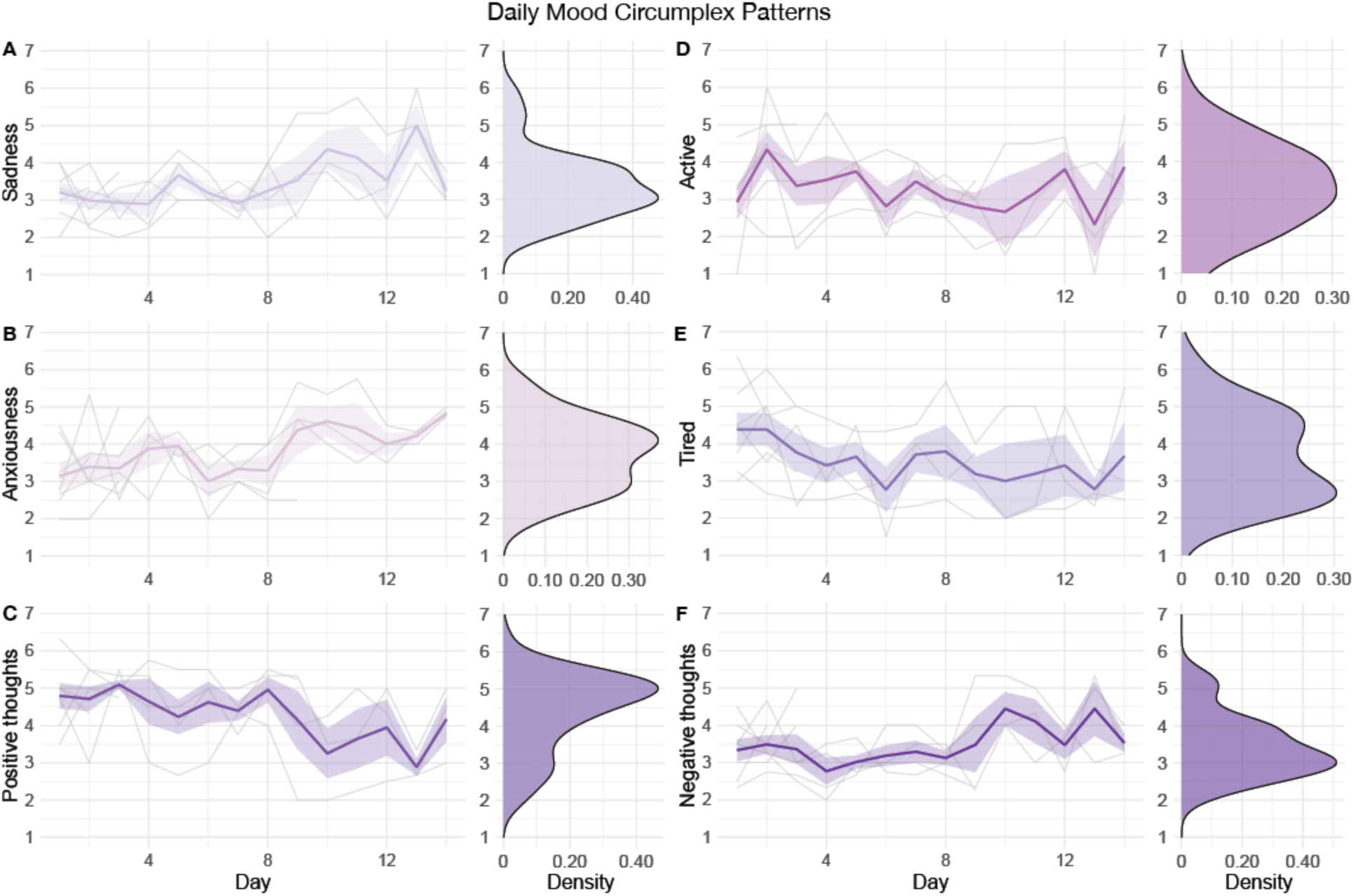
Ecological momentary assessment (EMA) daily mood patterns. Daily patterns for each participant across the 14-day EMA period are shown for mood circumplex items: (**A**) sadness, (**B**) anxiousness, (**C**) positive thoughts, (**D**) active, (**E**) tired, and (**F**) negative thoughts). For each mood circumplex item, the left-side figure displays the score of 1 through 7 for each of the 14 days, with individual trajectories (gray lines), the mean group trajectory (colored line), and standard errors (colored shade). The right-side figure displays the distribution density of each score for that mood circumplex item across all days and participants.

### Actigraphy

A sample of n = 6 participants completed the 21-day actigraphy procedure; actigraphy devices were not available for the first four participants enrolled in the study. We examined participants’ 24-hour sleep timing patterns through two measures: sleep onset and wake up (**Figure 7**). The preliminary data revealed differences in monophasic (i.e., single peak in the density plot of sleep onset) and biphasic (i.e., two peaks in the density plot of sleep onset) sleep patterns between participants. The occurrence of multiple peaks in a participant’s density plot suggests irregularity in sleep onset and wake up times. In addition to 24-hour sleep timing patterns, we explored daily sleep and physical activity patterns over multiple days and nights (**Figure 8**). Daily sleep metrics included sleep duration, wake after sleep onset (WASO; time spent awake after sleep onset but before the official wake up time), and sleep efficiency (the percentage of time asleep while in bed). The data collected for the 21-day actigraphy period revealed a mean sleep duration of 6.78 hours (SD = 1.22), a mean wake after sleep onset period of 1.34 hours (SD = 0.73), and mean sleep efficiency of 82% across participants (SD = 6.68). The data also highlighted distinct levels of physical activity (e.g., light activity, moderate activity, and vigorous activity). Light activity refers to an acceleration greater than 40 milli-g (mg), moderate activity refers to an acceleration greater than 100 mg, and vigorous activity refers to an acceleration greater than 400 mg. The actigraphy data revealed an across subject average of mean light activity level of 150 minutes per day (SD = 41.6), a mean moderate activity level of 130 minutes per day (SD = 43.7), and a mean vigorous activity level of 4.79 minutes per day (SD = 6.31).

**Figure 7.**
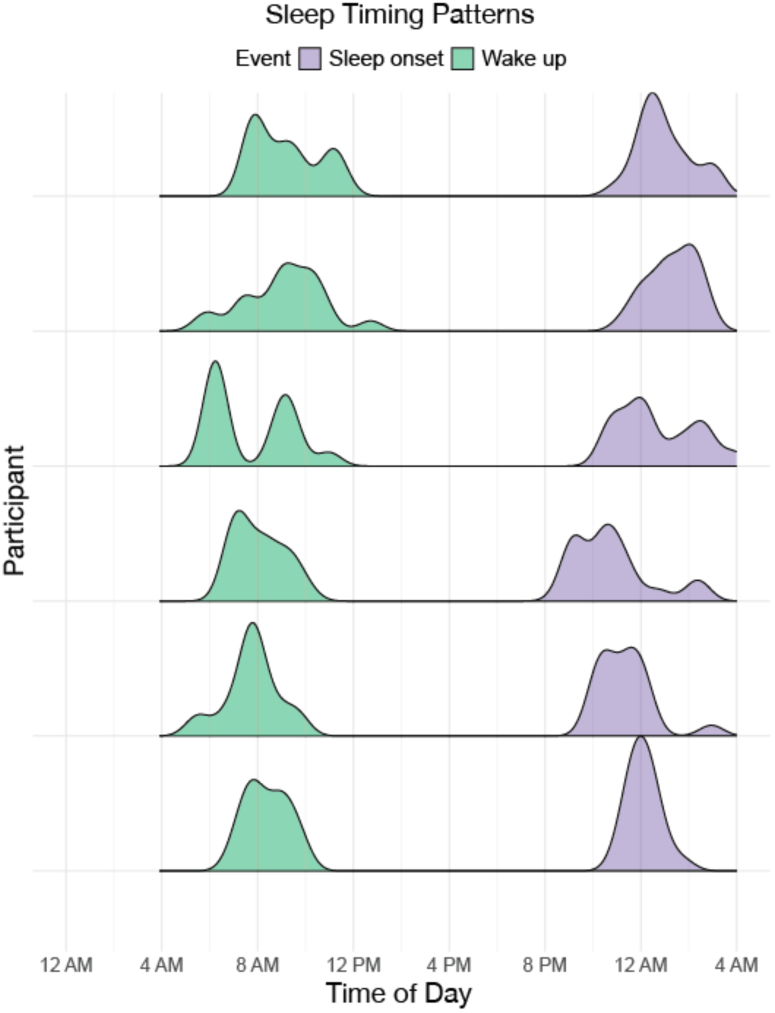
Sleep onset and wake timing. Distributions of sleep onset and wake times over a 24-hour period are displayed, with purple indicating sleep onset and green indicating wake up for each participant who completed the actigraphy procedure.

**Figure 8.**
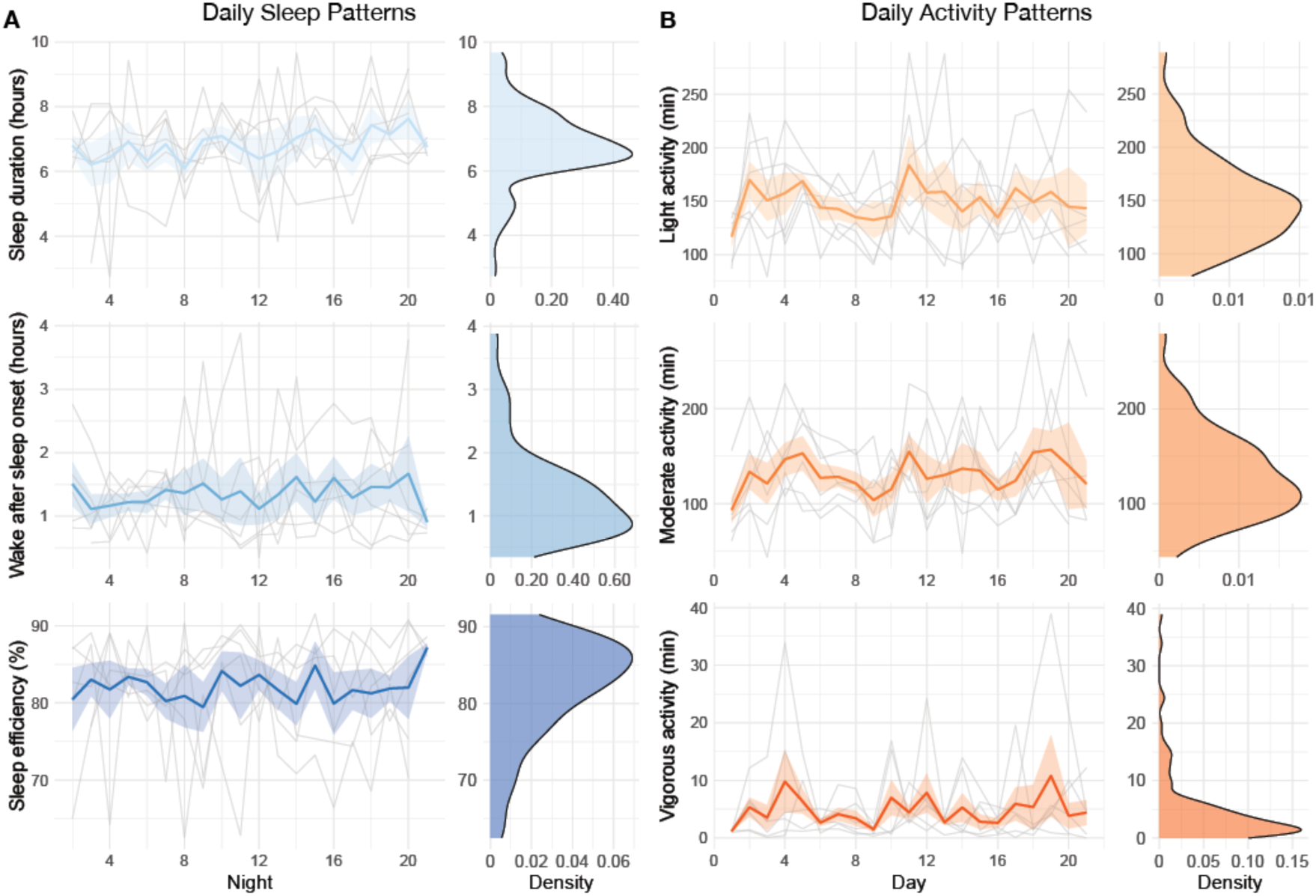
Daily sleep and physical activity patterns. Daily patterns for each participant across the 21-day actigraphy period are shown for: (**A**) daily sleep patterns (sleep duration, wake after sleep onset, and sleep efficiency) and (**B**) daily activity patterns (light activity, moderate activity, and vigorous activity). For each actigraphy measure, the left-side figure displays the measure across the 21 days, with individual trajectories (gray lines), the mean group trajectory (colored line), and standard errors (colored shade). The right-side figure displays the distribution density of the actigraphy measure.

### Neuroimaging

Neuroimaging data was collected on all n = 10 participants. Preprocessed structural MRI data exhibited expected mean cortical thickness, curvature, and sulcal depth across the sample (**Figure 9**). In addition, cerebral blood flow estimates from the ASL images displayed expected patterns of perfusion, with greater cerebral blood flow in cortical gray matter than white matter (**Figure 10**). For the multi-echo fMRI data, as expected, T2* did not change dramatically by including or excluding NORDIC denoising (**Supplementary Figure 1**). Notably, tedana and AROMA showed relatively high levels of agreement on the classification of noise components (**Supplementary Figure 2A**). However, tedana appeared to be more sensitive, identifying some noise components that AROMA did not. In contrast, AROMA classified very few components as noise that tedana did not identify. The noise components only identified by tedana explained somewhat lower variance than the components that both methods identified as noise (**Supplementary Figure 2B**). Mean correlation matrices for the “Bao” and “Your Friend the Rat” video watching tasks derived from fully processed fMRI data revealed expected patterns of functional connectivity across all runs (**Figure 11**). Finally, diffusion imaging data collected using CS-DSI displayed expected patterns of generalized fractional anisotropy (GFA) and mean diffusivity (MD), with GFA being higher in white matter and MD being higher in gray matter (**Figure 12**). Reconstructed white matter bundles in three exemplar participants revealed excellent delineation of individual-specific white matter tracts **(Figure 12C)**.

**Figure 9.**
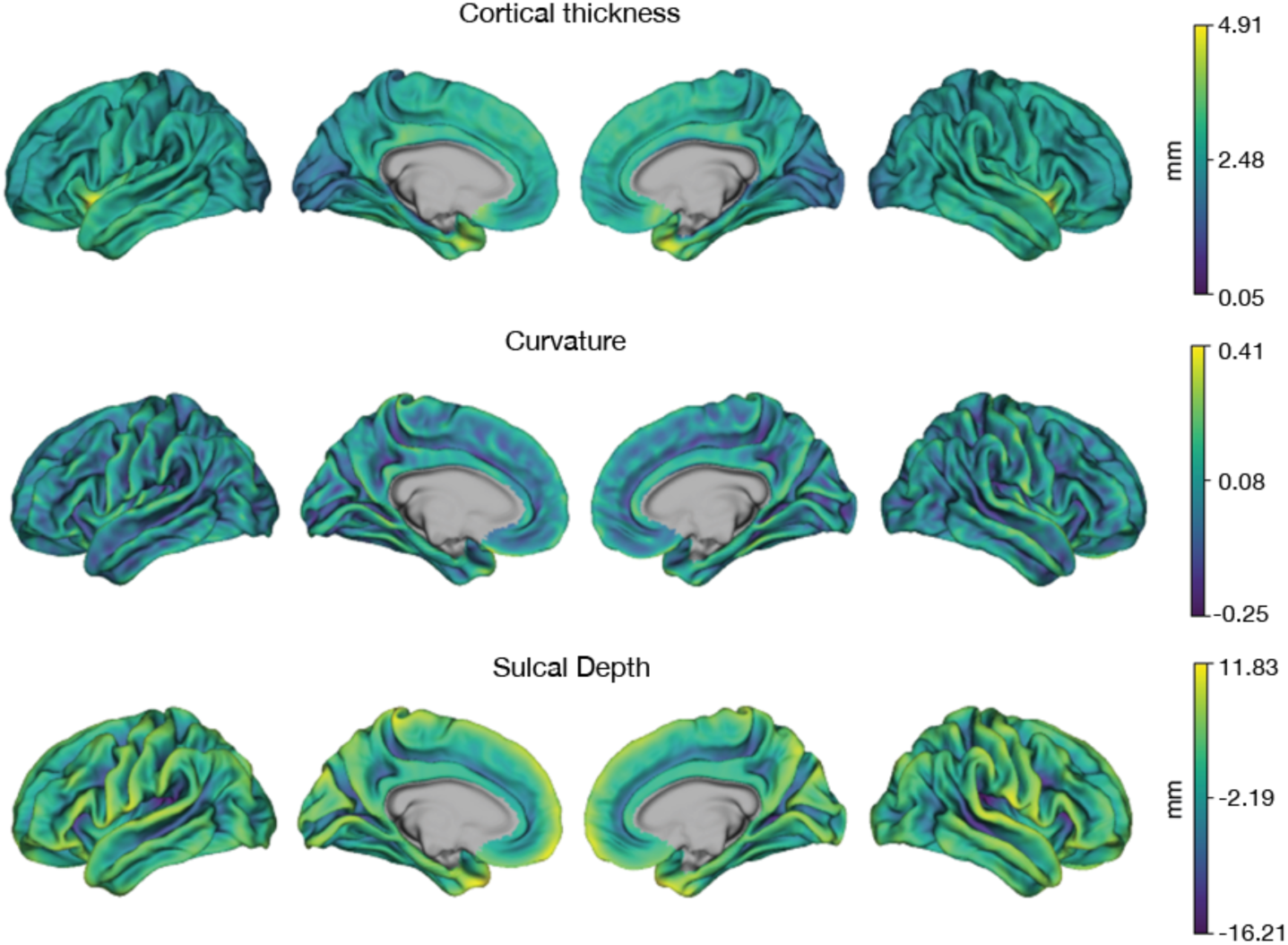
Anatomical MRI. Mean cortical thickness, curvature, and sulcal depth across the initial sample are displayed.

**Figure 10.**
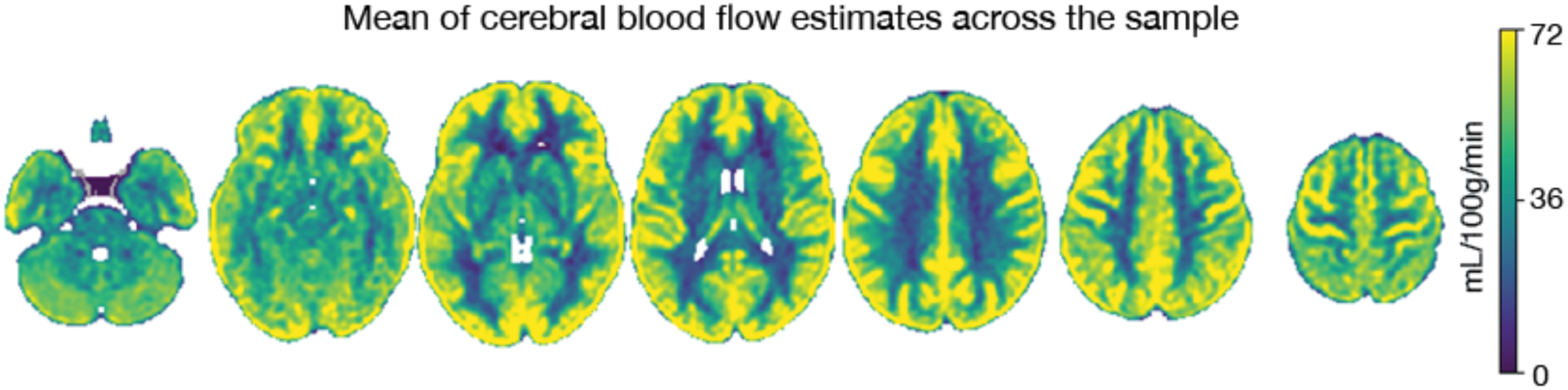
ASL cerebral blood flow. The mean of cerebral blood flow estimates across the initial sample is displayed.

**Figure 11.**
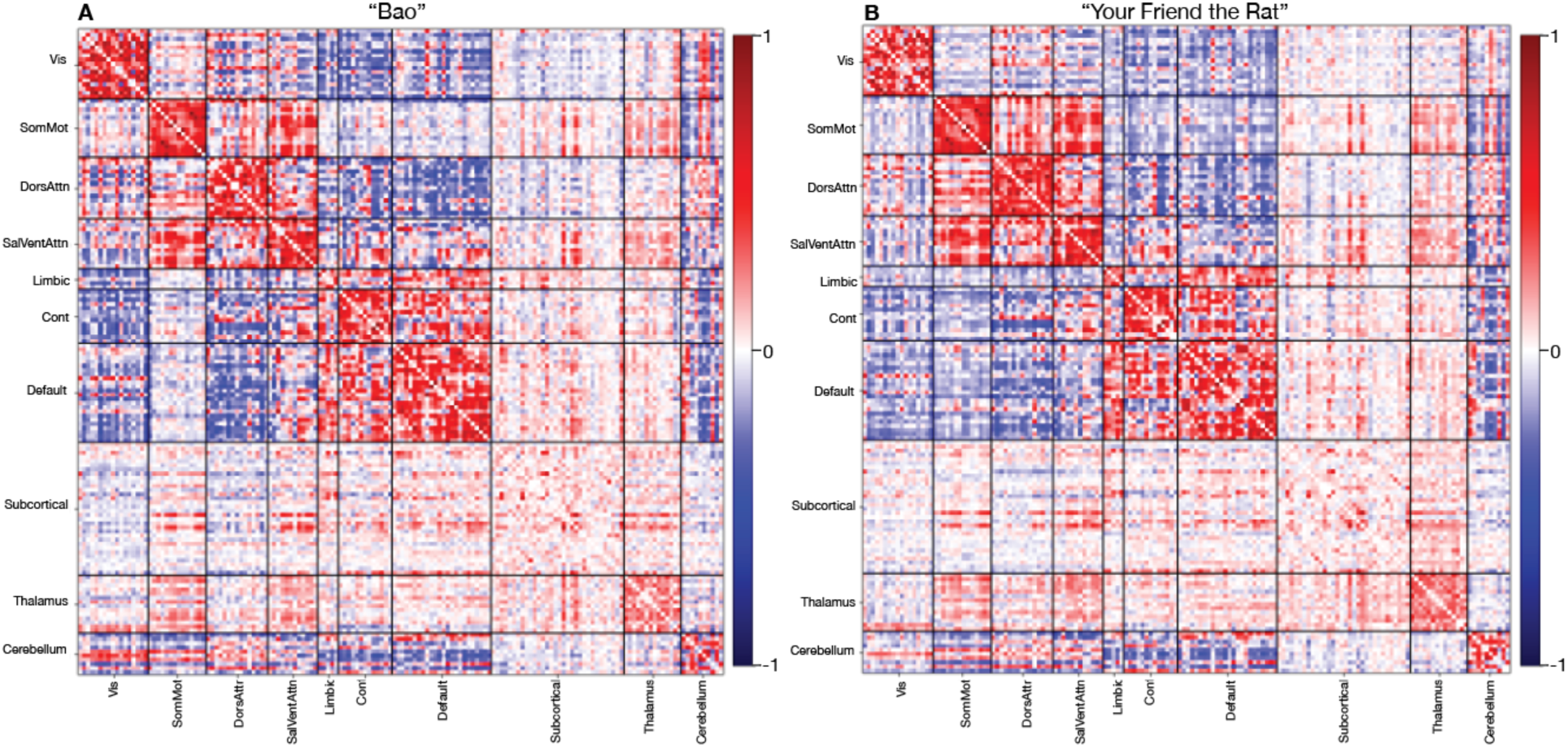
Multi-echo fMRI. The mean correlation matrices from (**A**) “Bao” and (**B**) “Your Friend the Rat” videos following complete preprocessing are displayed.

**Figure 12.**
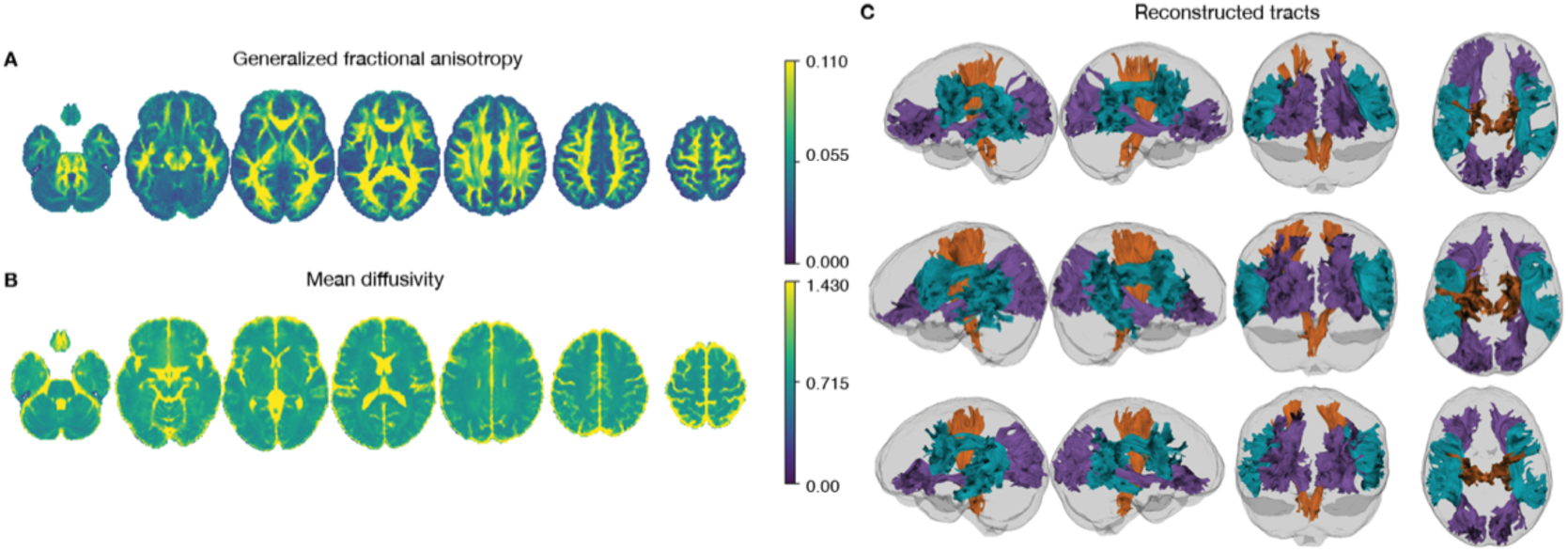
Compressed sensing diffusion spectrum MRI. Average generalized fractional anisotropy (**A**) and mean diffusivity (**B)** from the initial sample are displayed. Examples of reconstructed white matter tracts from three participants (participants 24683, 24053, and 60295) are displayed in C.

## Discussion

This study aims to provide high-quality brain and behavioral data to investigate affective dynamics in youth. Specifically, measures include advanced multimodal imaging, ecological momentary assessment (EMA), retrospective self-report questionnaires, computerized cognitive assessments, and actigraphy measures. Below we briefly discuss relevant context, limitations, and important future directions.

Difficulties in affective regulation during childhood may confer increased risk for psychopathology in adulthood. Children rely on social interactions to regulate their mood by seeking reassurance from others, eventually developing an internal self-regulation mechanism to manage their affective state.^1^ Deficits in the child-caregiver relationship and early exposure to trauma may contribute to the development of affective instability.^1,173^ Affective instability involves acute fluctuations in affect and is linked to aberrant development of psychological and psychosocial capabilities (i.e., shortfalls in self-esteem, social interactions, and sense of identity) and increased vulnerability to psychopathology later on.^1,174^

While affective instability is present in multiple psychiatric disorders, it has typically been studied within a single diagnosis using a case-control design, limiting the ability to examine common brain circuits impacted across diagnoses.^175,176^ Affective instability is not only a diagnostic criterion of borderline personality disorder (BPD)^177^ but is also observed in diverse conditions seen among adolescents and young adults, including bipolar disorder,^2^ major depressive disorder (MDD),^2^ and attention-deficit hyperactivity disorder (ADHD).^2^ Existing evidence suggests that affective instability is associated with suicide threats and attempts among participants with BPD^4^ as well as suicidal ideation in adults with other psychiatric disorders (e.g., depression, anxiety).^178^ To extend prior work studying affective instability beyond traditional diagnostic categories, this study characterizes affective instability through a transdiagnostic, dimensional approach. Dimensional approaches to studying psychopathology have the potential to enhance the understanding of how maladaptive behaviors gradually evolve across development.^176^ By recruiting a community sample of youth and collecting moment-to-moment information on affective dynamics, passive physical activity and sleep data, and high-quality neuroimaging data, the data from this study may accelerate research linking brain circuits to affective instability as a transdiagnostic construct.

In addition to densely sampled behavioral and clinical measures, this study leverages cutting-edge neuroimaging methods -- such as multi-echo fMRI -- to examine the neural substrates of affective dynamics. As a result of small effect sizes, previous neuroimaging work on mental illness has often required very large sample sizes to reliably identify relationships between brain features and behavioral phenotypes.^39,41,179,180^ However, by adopting measures to enhance reliability and detection of individual-specific variation, studies may potentially mitigate the need to rely on large samples to identify meaningful brain-behavior associations.^44^ One promising approach is using personalized mapping of individuals’ functional neuroanatomy, which has been shown to better predict behavioral phenotypes than group average parcellations.^181,182^ Personalized functional networks (PFNs) account for individual variation in brain organization and have the potential to enhance the sensitivity of fMRI data to diagnosis-specific variation.^182,183^ For example, expanded topography of the frontostriatal salience network defined using PFNs has been shown in individuals with depression.^184^ In developmental cohorts, PFN topography is refined through childhood and adolescence and exhibits prominent sex differences.^185,186^ These findings highlight the potential for studying PFNs in a youth population. As multi-echo fMRI allows PFNs to be identified with smaller amounts of neuroimaging data, an advance that significantly improves the feasibility of scanning youth populations,^48^ we anticipate that the multi-echo fMRI data generated from this study will allow accurate delineation of PFNs. When coupled with densely-sampled behavioral EMA and actigraphy data, we hope that multi-echo fMRI will enhance our ability to detect brain-behavior relationships relevant for affective instability.

Although the current study introduces important advances, several limitations should be acknowledged. The study aims to capture the developmental relevance of affective instability through a community sample of 13- to 23-year-old participants. However, participants are only measured at one time point, precluding inference on within-individual development. Furthermore, because the current participant pool is collected through community sampling, and participants do not complete a diagnostic interview, the study does not examine specific clinical diagnoses. Finally, our target sample is only 100 individuals; such a sample may still have limited statistical power to find brain-behavior links despite advances in behavioral assessment (e.g., EMA) and neuroimaging methods (e.g., multi-echo fMRI).

Taken together, this study provides a novel data resource to examine affective dynamics using precision brain and behavioral approaches – including EMA sampling, actigraphy data collection, and multi-echo fMRI. Moving forward, future longitudinal studies of brain development that examine participants pre- and post-adolescence may more fully characterize the developmental progression of affective instability, especially in clinical populations. We hope that this data resource will contribute to the identification of early markers of affective instability and emotion dysregulation before they develop into more severe psychopathology.

## Data availability

All raw and processed imaging data is available on OpenNeuro: Raw https://openneuro.org/datasets/ds006131; MRIQC https://openneuro.org/datasets/ds006143; QSIPrep https://openneuro.org/datasets/ds006182; QSIRecon https://openneuro.org/datasets/ds006184; fMRIPrep https://openneuro.org/datasets/ds006185; ASLPrep https://openneuro.org/datasets/ds006188; fMRIPost-AROMA https://openneuro.org/datasets/ds006189; tedana https://openneuro.org/datasets/ds006190; tedana & AROMA https://openneuro.org/datasets/ds006191; XCP-D https://openneuro.org/datasets/ds006192 }.

## Code availability

All analysis code for this project is openly available on GitHub: https://github.com/PennLINC/affective-instability.

## Acknowledgments

This study was supported by the AE Foundation, The Dean’s Innovation Fund at the University of Pennsylvania Perelman School of Medicine, and the Penn/CHOP Lifespan Brain Institute. Additional support was provided by grants from the National Institute of Health: R01MH113550 (T.D.S.), R01MH112847 (R.T.S.), R01EB022573 (T.D.S.), R37MH125829 (T.D.S & D.A.F.), R01MH117014 and R01MH119219 (R.C.G., R.E.G.), K23 MH133118 (E.B.B.), BBRF #31319 (E.B.B.), R01EB031080 (J.A.D., M.D.T, & M.T.), BBRF #30837 (S.S.), BWF 1022955 CAMS (S.S.), DP5OD036142 (S.S.), U24NS130411 (D.W.), S10OD023495 (D.W.), F31MH136685 (J.B.).

## Conflicts of interest

**MT is employed by Siemens Medical Solutions USA**

## Supplemental Tables

**Supplementary Table 1.**
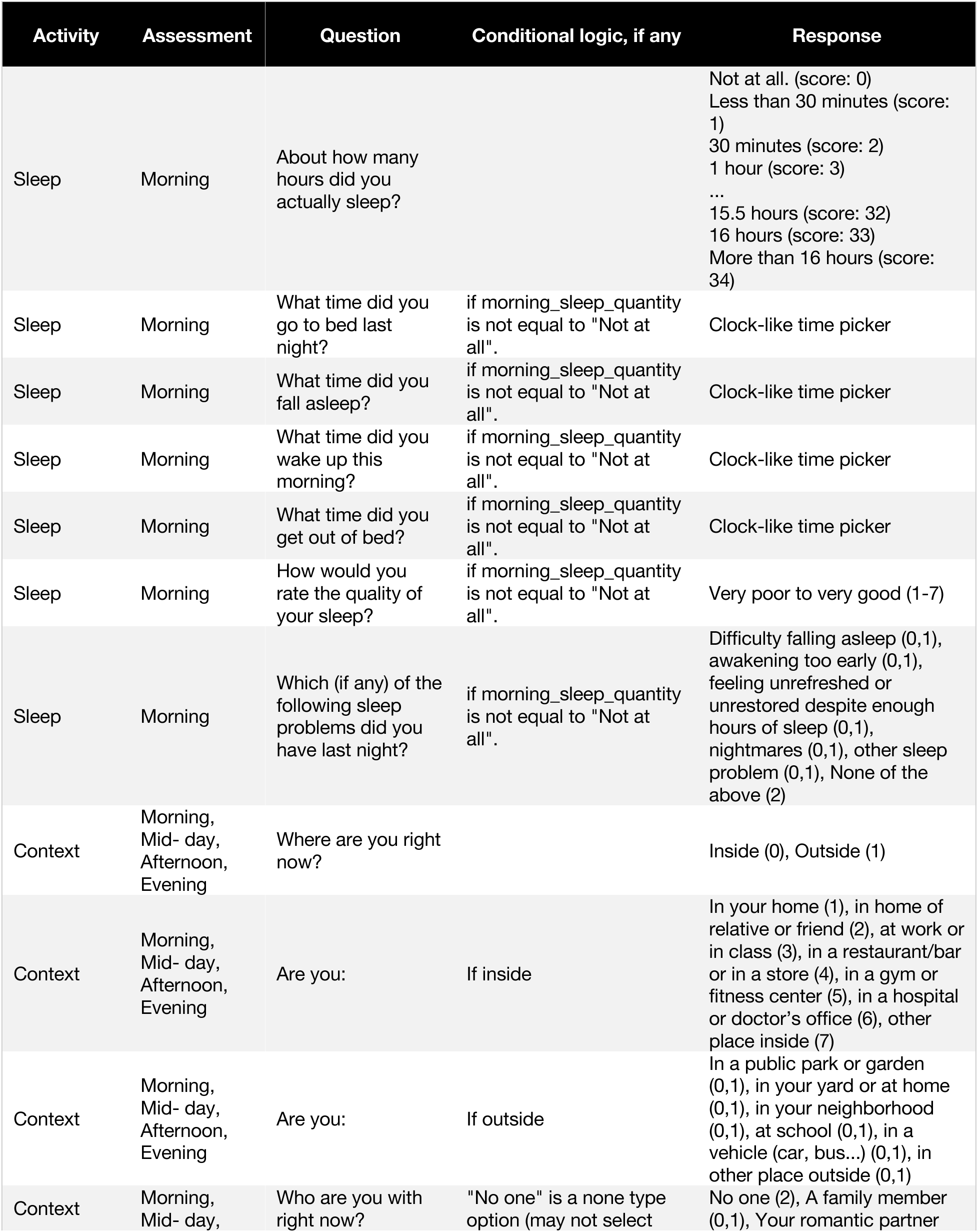
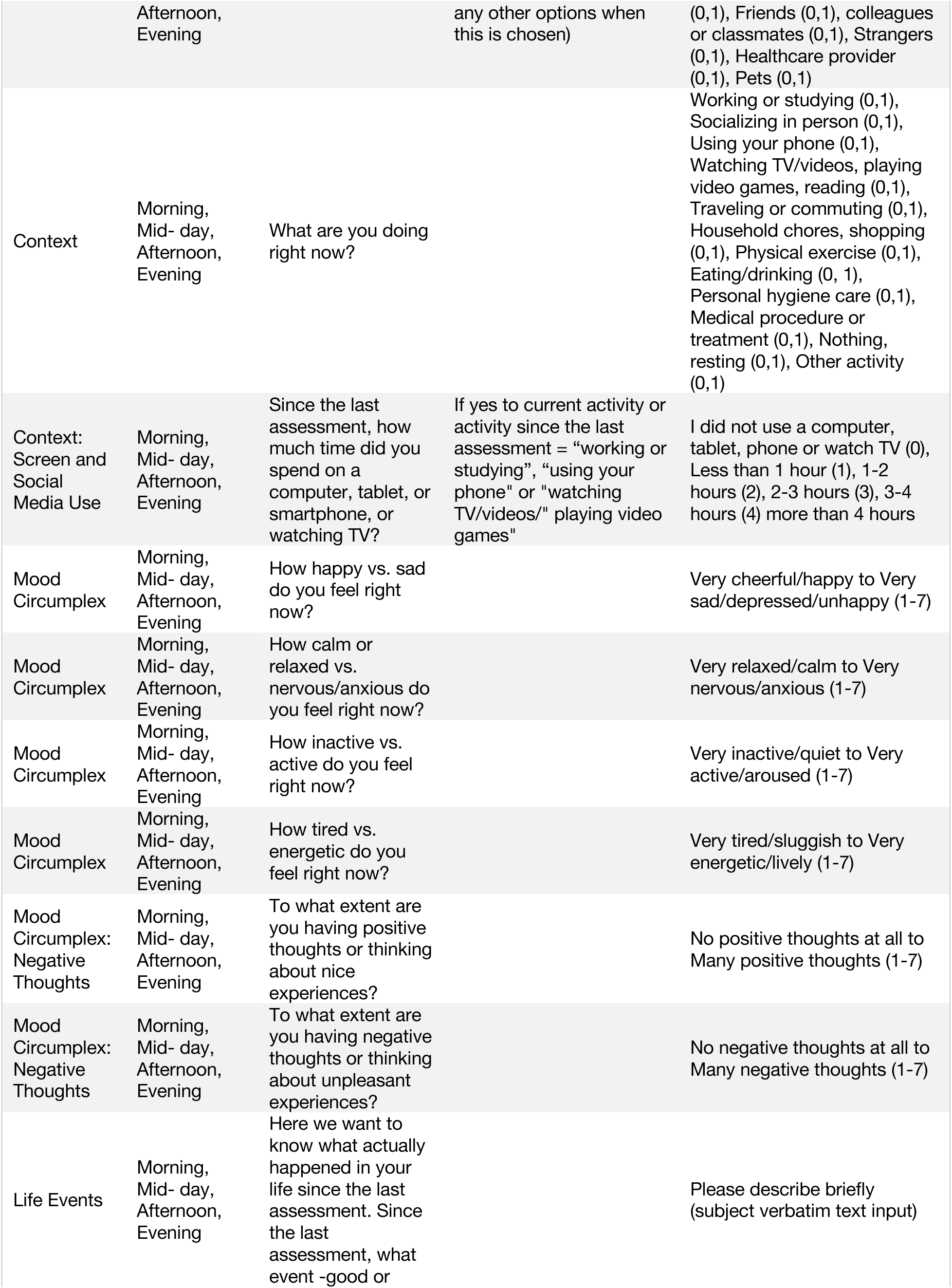
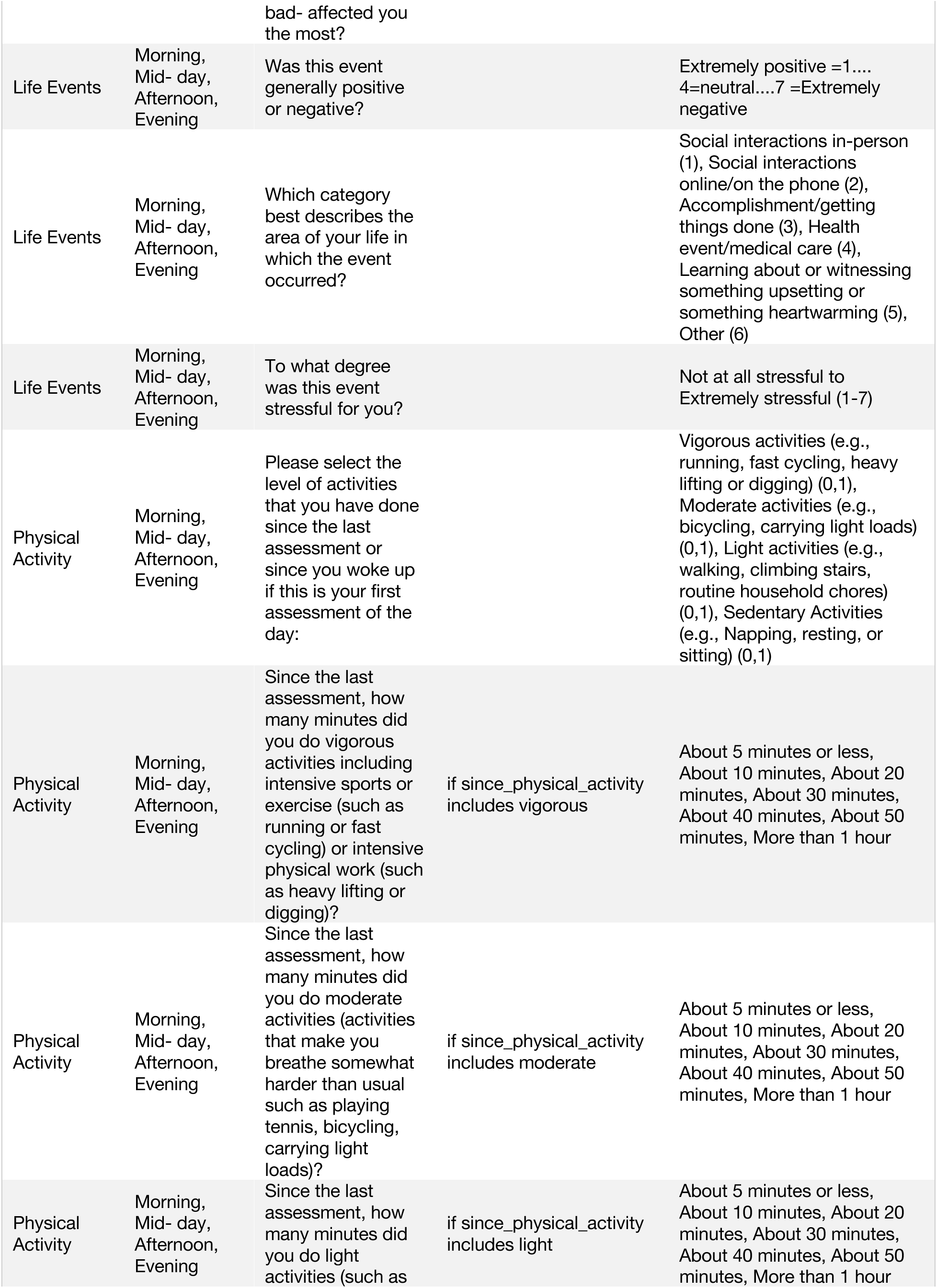
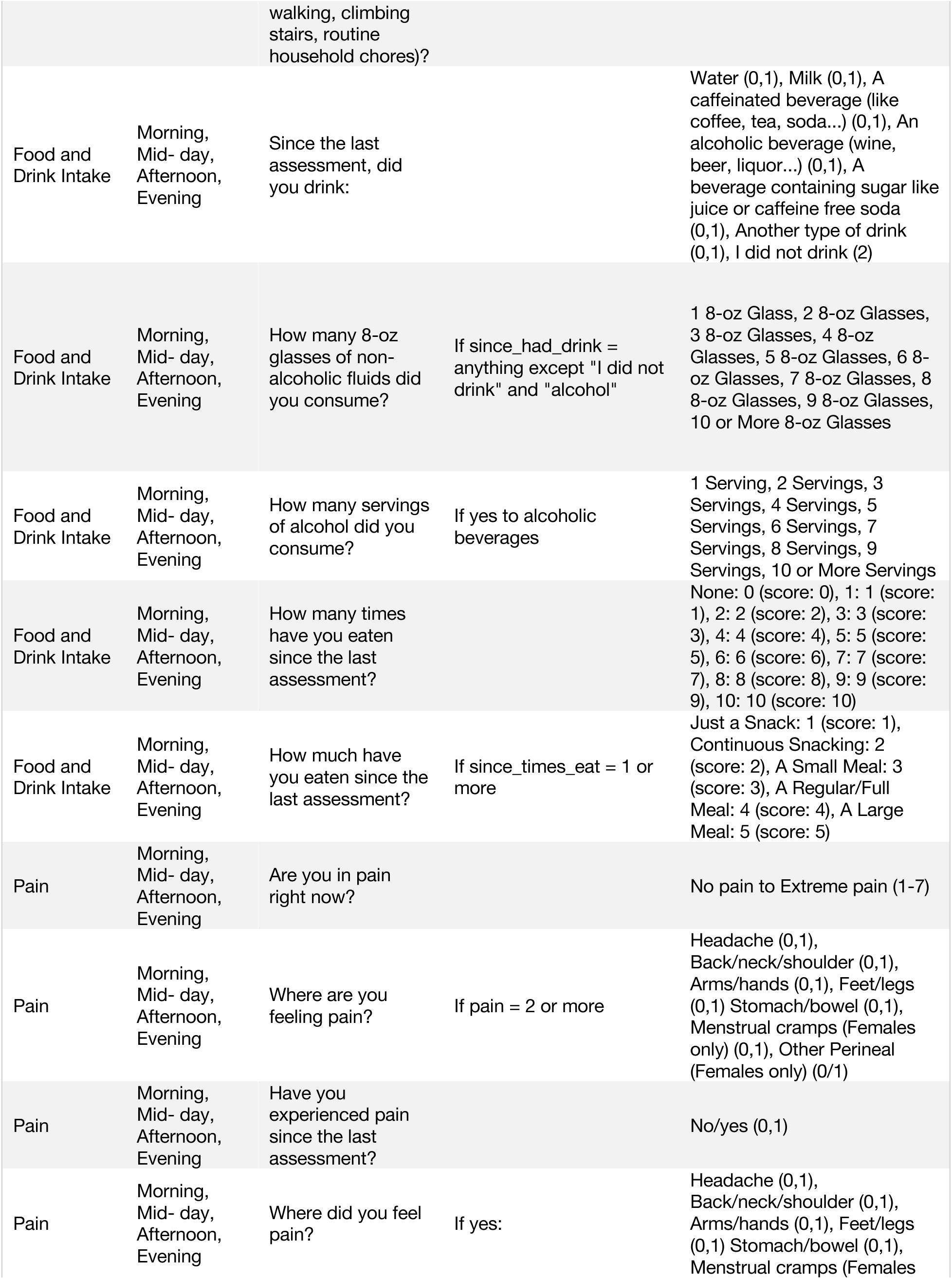
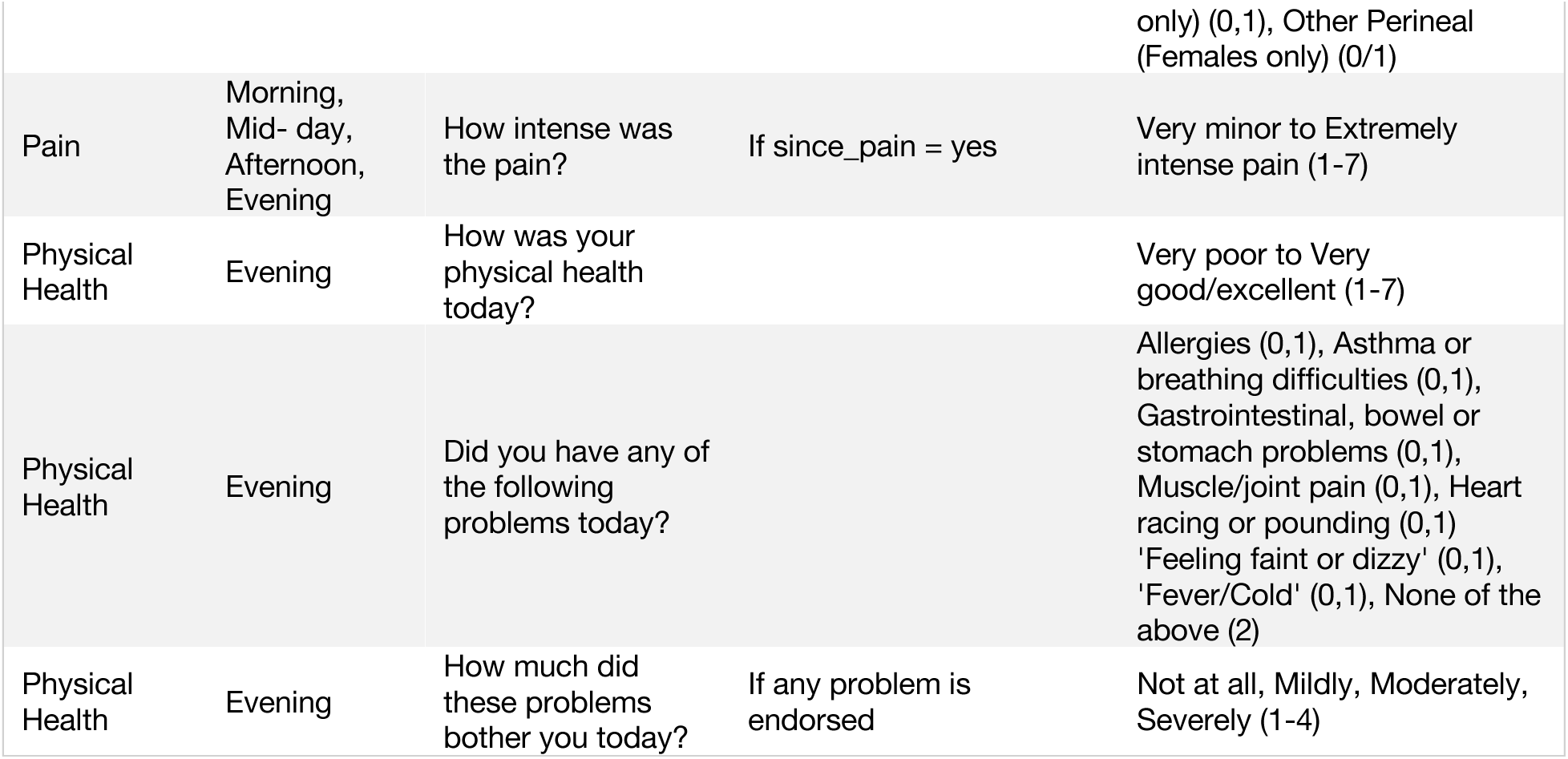
Ecological momentary assessment (EMA) items.

## Supplemental Figures

**Supplementary Figure 1.**
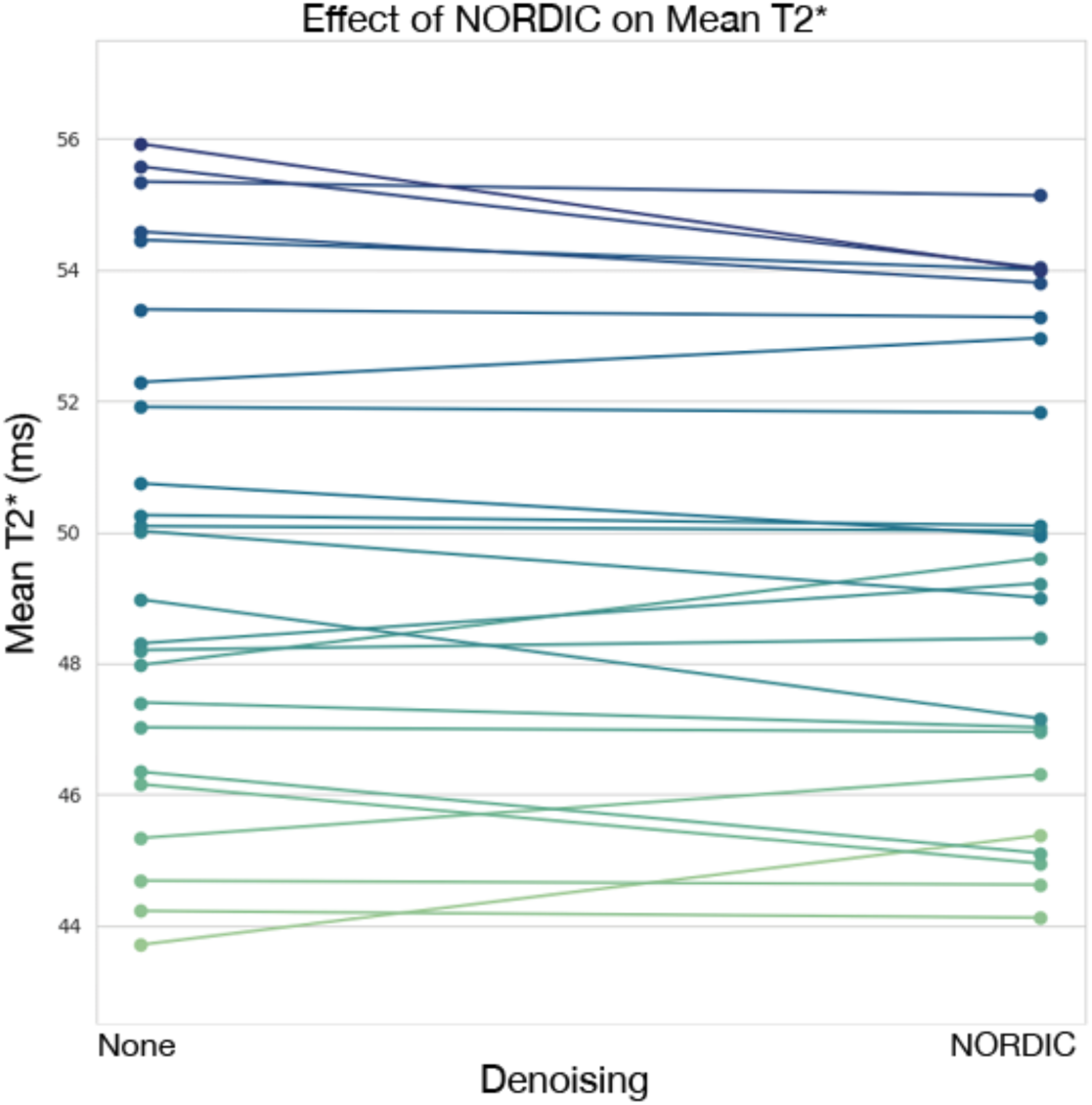
Effect of NORDIC denoising on mean T2*. The effect of NORDIC denoising on mean T2* found in preprocessed multi-echo fMRI is displayed. The inclusion and exclusion of NORDIC denoising did not dramatically change T2*.

**Supplementary Figure 2.**
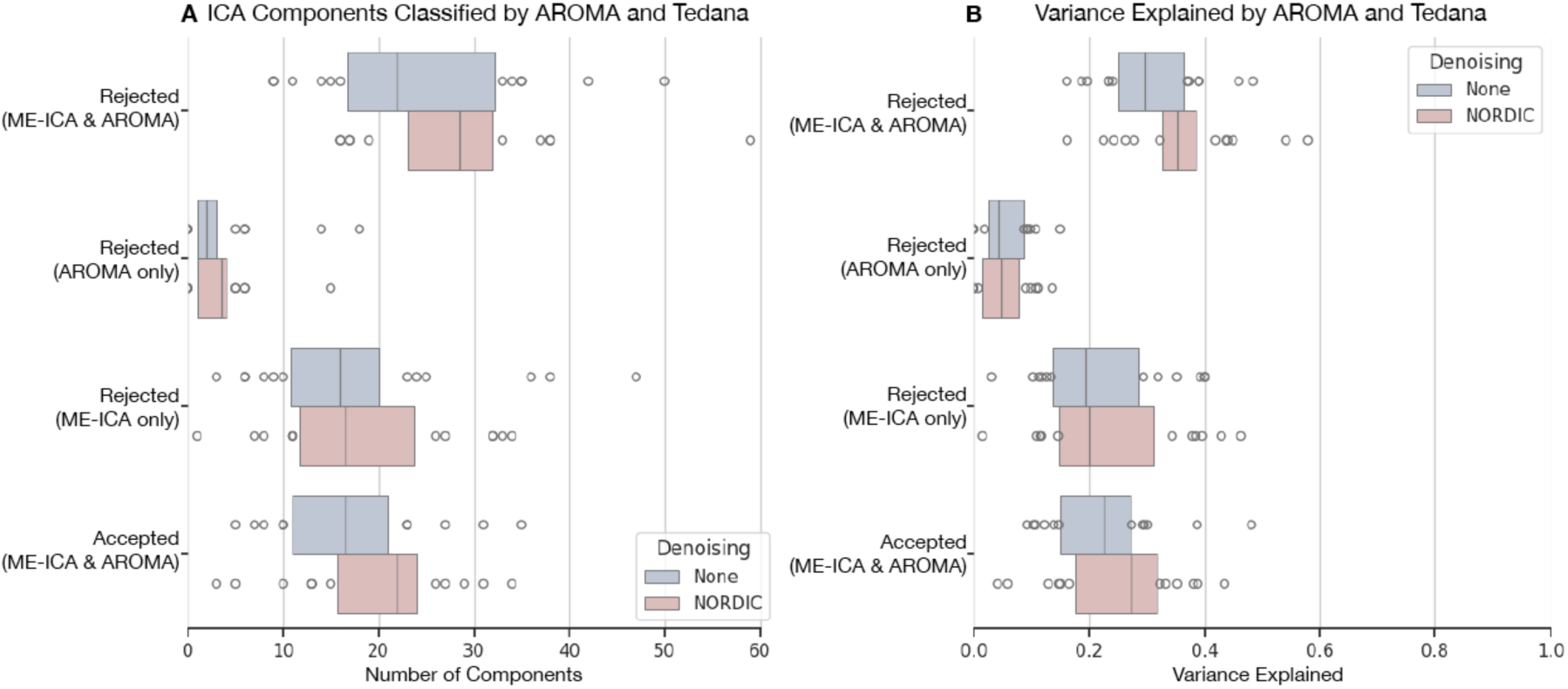
Tedana and AROMA noise component classifications. Classification of noise components by tedana, which implements multi-echo ICA (ME-ICA), and AROMA are displayed. **A)** A plot of ICA components identified by tedana and AROMA shows that tedana identifies some noise components that AROMA does not. AROMA classifies very few components as noise that tedana does not identify as such. **B)** Comparing the amount of variance in the data attributed to these noise ICA components, tedana accounts for a larger proportion of variance as noise compared to AROMA. The blue and red bars show the effect of no NORDIC versus with NORDIC denoising, respectively.

## Notes

https://openneuro.org/datasets/ds00613

https://openneuro.org/datasets/ds006143

https://openneuro.org/datasets/ds006182

https://openneuro.org/datasets/ds006184

https://openneuro.org/datasets/ds006185

https://openneuro.org/datasets/ds006188

https://openneuro.org/datasets/ds006189

https://openneuro.org/datasets/ds006190

https://openneuro.org/datasets/ds006191

https://openneuro.org/datasets/ds006192

### Competing Interest Statement

Manuel Taso is employed by Siemens Medical Solutions USA

### Summary of Updates

Correcting a typo in the abstract of the main manuscript file.

